# Noradrenergic Tone is Not Required for Neuronal Activity-Induced Rebound Sleep in Zebrafish

**DOI:** 10.1101/2023.03.17.533174

**Authors:** Eleanor Benoit, Declan G Lyons, Jason Rihel

**Affiliations:** Department of Cell and Developmental Biology, University College London, WC1E 6BT, United Kingdom

**Author notes:** Corresponding author: Jason Rihel.

**Keywords:** *c-fos*, noradrenalin, sleep, sleep homeostasis, zebrafish

## Abstract

Sleep pressure builds during wakefulness, but the mechanisms underlying this homeostatic process are poorly understood. One zebrafish model suggests that sleep pressure increases as a function of global neuronal activity, such as during sleep deprivation or acute exposure to drugs that induce widespread brain activation. Given that the arousal-promoting noradrenergic system is important for maintaining heightened neuronal activity during wakefulness, we hypothesised that genetic and pharmacological reduction of noradrenergic tone during drug-induced neuronal activation would dampen subsequent rebound sleep in zebrafish larvae. Unexpectedly, dampening noradrenergic tone with the α_2_-adrenoceptor agonist clonidine during acute caffeine or pentylenetetrazol treatment enhanced subsequent rebound sleep, while stimulating noradrenergic signalling during caffeine exposure with a cocktail of α_1_- and β-adrenoceptor agonists did not enhance sleep. Similarly, CRISPR/Cas9-mediated elimination of the *dopamine β-hydroxylase* (*dbh*) gene, which encodes an enzyme required for noradrenalin synthesis, enhanced baseline sleep in larvae but did not prevent additional rebound sleep following acute induction of neuronal activity. Across all drug conditions, *c-fos* expression immediately after drug exposure varied inversely with noradrenergic tone and correlated strongly with the amount of induced rebound sleep. These results are consistent with a model in which increases in neuronal activity, as reflected by brain-wide levels of *c-fos* induction, drive a sleep pressure signal that promotes rebound sleep independently of noradrenergic tone.

## Introduction

Sleep is a widespread – possibly universal – feature of animal life (Keene and Duboue, 2018), but its definitive purposes continue to elude us. There is increasing acknowledgement, however, that the functions of sleep relate primarily to the brain (Hobson, 2005), perhaps encompassing the replenishment of cerebral energy stores depleted during waking (Benington and Heller, 1995) and memory consolidation (Rasch and Born, 2013). The timing, duration and intensity of sleep are regulated per the “two-process” model, in which an animal’s circadian rhythm dictates the time(s) of day when it will tend to sleep, while homeostatic sleep pressure accumulates during waking to drive changes in the depth and duration of sleep (Borbély and Achermann, 1999). How and where homeostatic sleep pressure accumulates as a function of brain-related processes remains poorly understood.

One idea is that homeostatic sleep need reflects the overall level of brain activity integrated over prior waking. While sleep pressure has traditionally been associated with wake duration (Borbély and Achermann, 1999), not all waking behaviour involves equivalent neuronal activity (Fisher et al., 2016; Milinski et al., 2021) and within-waking arousal states can modulate the accumulation of sleep pressure (Yamagata et al., 2021; Vassalli and Franken, 2017). Experiments in zebrafish have demonstrated that acutely and transiently elevating neuronal activity with arousing drugs such as caffeine is followed by increased sleep (Reichert et al., 2019). This drug-induced rebound sleep is dissociable from prior wake time and physical hyperactivity but correlates strongly with the level of preceding global neuronal activity as measured by *c-fos* expression and whole-brain calcium imaging. Consistent with this, the intensity of regional neuronal activity during waking in mammals is associated with the extent of local offline periods and changes in regional slow-wave activity (a measure of sleep pressure) during the following sleep period (Krueger et al., 2019), while in mice, global slow-wave activity during NREMS has been shown to reflect the integrated cortical neuronal activity levels of the preceding wake period (Thomas et al., 2020). How widespread changes in neuronal activity would ultimately trigger changes in whole animal sleep is unclear, but evidence in both mice (Ma et al., 2019) and zebrafish (Reichert et al., 2019) implicates galaninergic neurons of the anterior hypothalamus and preoptic area (POA) as an effector arm of homeostatic sleep regulation.

One vital system for maintaining brain-wide arousal and implicated in *c-fos* expression during waking is the noradrenergic system (Cirelli and Tononi, 2000). The locus coeruleus (LC) is a small neuronal population (∼10-20 neurons in zebrafish; Farrar et al., 2018) that is the chief source of noradrenalin in the brain (Chandler et al., 2019) and is highly conserved among vertebrates, including zebrafish (Wang et al., 2022). LC neurons ramify widely, such that noradrenalin can act throughout the brain (Du et al., 2018) and also inhibit sleep-active neurons of the POA (Liang et al., 2021; Nelson et al., 2003). Indeed, the activity of the LC is intimately coupled with the sleep/wake behavioural state of the animal, and noradrenergic signalling is required for the normal maintenance of the waking state in animals including mice and zebrafish larvae (Ouyang et al., 2004; Singh et al., 2015). During waking, the LC is tonically active; this activity falls substantially during non-REM sleep (NREMS) (Steininger et al., 2001) and virtually ceases during REM sleep (Jones, 1991). Activity in the LC precedes spontaneous waking (Saper et al., 2010), and activation of the LC during sleep can cause immediate sleep-to-wake transitions (Carter et al., 2010). Additionally, phasic burst firing of the LC in response to a salient stimulus (Carter et al., 2010) helps the animal focus its attention (Jones, 1991). As such, the maintenance of brain-wide noradrenergic tone is thought to be crucial to sustaining wake-related arousal and neuronal activity, and is a candidate driver of sleep need (Cirelli et al., 2005).

Here, we explore the role of the noradrenergic system in modulating stimulant drug-induced sleep pressure in zebrafish larvae. Genetic and pharmacological manipulation of noradrenergic transmission surprisingly reveals that lowered noradrenergic tone enhances both stimulant-drug-induced *c-fos* induction and subsequent rebound sleep. This presents a new insight into the relationship of noradrenergic activity and sleep pressure generation and is consistent with a model whereby increases in neuronal activity, as reflected by *c-fos* expression, can generate homeostatic sleep drive independently of the noradrenergic system.

## Methods and materials

All animal protocols were performed in accordance with project licence PA8D4D0E5, awarded to Jason Rihel by the UK Home Office under the UK Animals (Scientific Procedures) Act 1986. Experiments used AB/Tupfel long-fin larvae up to 8 days post fertilisation (dpf), before the onset of sexual maturation.

### Sleep/wake activity assays

Embryos were reared in an incubator at 28.5°C on a 14hr:10hr light:dark cycle, with lights on from 9am (zeitgeber time zero = ZT0). At 5 dpf, individual larvae were pipetted into each well of a 96-square well plate (Whatman). Each well contained 650µl of fish water (0.3g/L Instant Ocean with 40µg/L of methylene blue). Wells were topped-up daily with fish water.

Videotracking was conducted per Reichert et al. (2019), using an automated Zebrabox system (ViewPoint Behaviour Technology) and maintaining a 14hr:10hr light:dark schedule. Ambient temperature was held at 26-28.5°C. Constant infrared illumination allowed for videotracking throughout the day/night cycle. “Quantization mode” in the ZebraLab software was used to record larval movements (detection parameters: sensitivity 20, burst 200, freeze 3 and bin size 60s). Custom “sleep_analysis2020” and “sleep_analysis_widget” MATLAB (MathWorks) codes were used to analyse the Zebrabox activity data (available on GitHub, https://doi.org/10.5281/zenodo.7644073). Sleep was identified as periods of inactivity lasting ≥ 1min, as such quiescent bouts have been shown to fulfil the criteria for a behavioural definition of sleep, including an elevated arousal threshold (Prober et al., 2006).

To pharmacologically compromise noradrenergic signalling, the α_2_-adrenoceptor agonist clonidine was added to the fish water on the afternoon of 5 dpf. A 1mM working solution of clonidine was prepared in 10% dimethyl sulfoxide (DMSO); 3.25µl of this was pipetted into each 650µl well to give a final concentration of 5µM clonidine (after Singh et al., 2015) and 0.05% DMSO. For control wells, 3.25µl of 10% DMSO was applied to give a final concentration of 0.05% DMSO.

To pharmacologically activate the noradrenergic system, a mixture of the α_1_-adrenoceptor agonist phenylephrine and the β- adrenoceptor agonist isoproterenol was added to the fish water from ZT0 + 10min at 6 dpf. A working solution of 0.5mM phenylephrine and 0.5mM isoproterenol was prepared in double distilled water. 13µl of this was pipetted into each 650µl well to give a final concentration for each drug of 10µM (after Yin et al. (2009), who found that either 10µM phenylephrine or 10µM isoproterenol alone significantly increased the zebrafish larval heart rate, and Rihel et al. (2010), who found that ∼10µM isoproterenol decreased larval sleep behaviour).

On 6 dpf at ZT1, the stimulant drugs caffeine or pentylenetetrazol (PTZ), or the same volume of water, were added to individual wells at 20s intervals. Caffeine, which antagonises adenosine-receptors (Porkka-Heiskanen and Kalinchuk, 2011), was applied at 2mM final concentration. PTZ, a GABA_A_-receptor antagonist, was applied at 10mM final concentration (see Key Resources table for working solution concentrations). After 1hr of caffeine/PTZ treatment, at ZT2, drug wash-off began. Each larva was individually pipetted into a 13.5cm diameter petri dish containing ∼150ml fish water, and then into a second 13.5cm water dish, and then into its respective well in a fresh 96-well plate. In Figs. 1, 4, 5 and 6, the blanked-out region on each sleep trace indicates this drug wash-off period, when the larvae were removed from the video tracking apparatus. The wash-off process took about 20s for each larva. Videotracking then resumed for two days and nights.

**Fig. 1.**
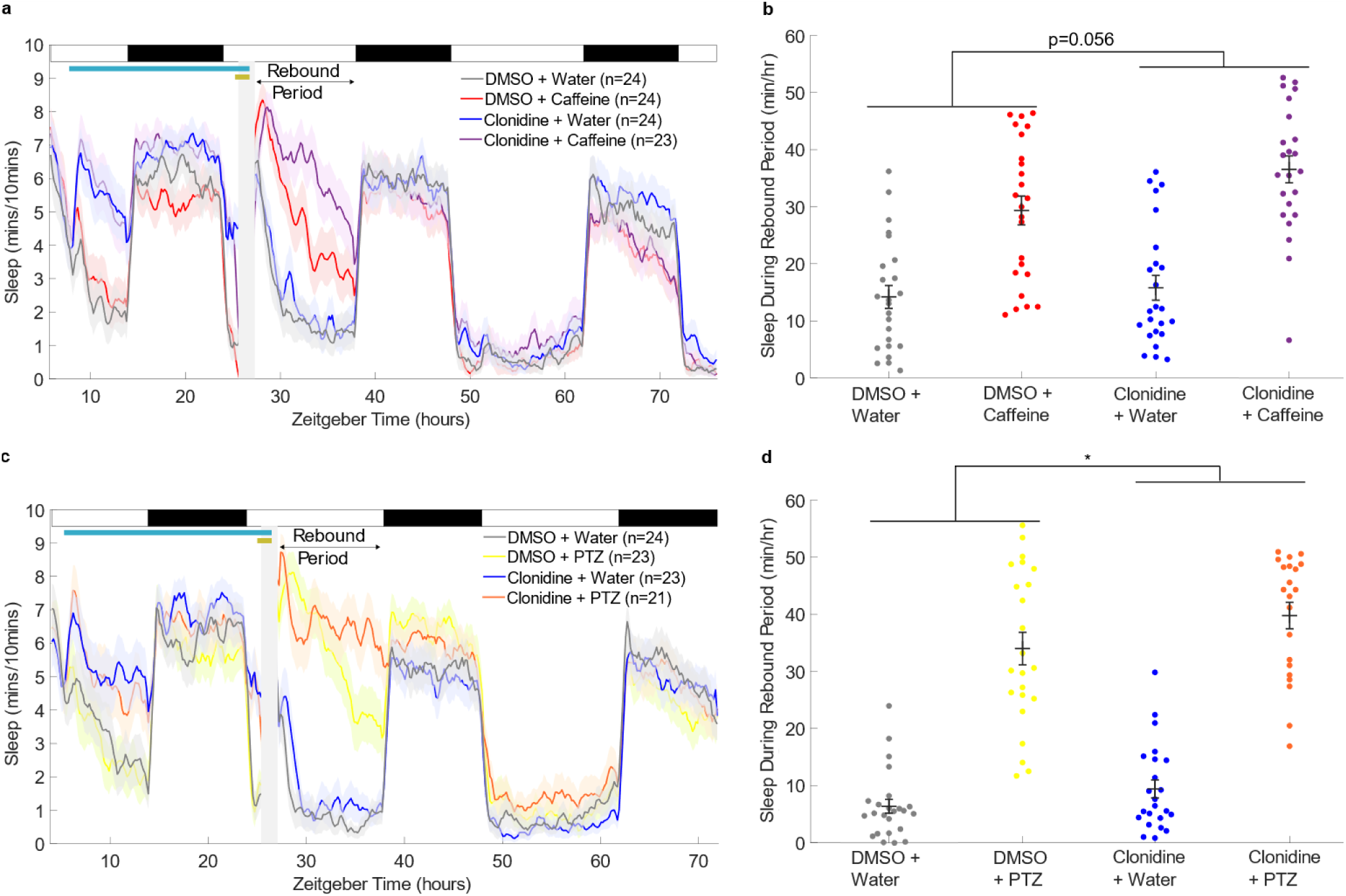
Activating α2-adrenoceptors during drug-induced arousal facilitates rebound sleep. **a** Sleep traces (± SEM) beginning at 5 dpf and continuing over three days and nights (time zero = first lights on) for larvae exposed to combinations of 5µM clonidine/DMSO and 2mM caffeine/water. Following drug wash-off, larvae experience rebound sleep (labelled Rebound Period). At the top, white and black bars represent day and night, respectively; the pale blue horizontal bar shows the clonidine exposure window, while the gold bar indicates the presence of stimulant. **b** shows the average total sleep/hr during the rebound period for each larva (black bar: mean ± SEM). Caffeine significantly increased rebound sleep (p=6.1 x 10^-12^, F(1,91) = 62.45), while clonidine trended to further enhance sleep (p=0.056, F (1,91) = 3.76, see also Fig. S1). There was no significant interaction between clonidine and caffeine treatment (p=0.22, F(1,91) = 1.52), based on a two-way ANOVA (caffeine treatment x clonidine treatment). **c** Sleep traces as in **a** for larvae exposed to combinations of clonidine and 10mM PTZ. The post-drug rebound sleep period of **c** is summarised for each larva in **d**. PTZ treatment significantly increased rebound sleep (p=4.9 x 10^-24^, F(1,87) = 196.55), as did clonidine treatment (p=0.037, F(1,87) = 4.48), while there was no significant interaction between clonidine and PTZ treatment (p=0.50, F(1,87) = 0.45), based on a two-way ANOVA (PTZ treatment x clonidine treatment). *p<0.05

### Drug treatment for quantitative real-time polymerase chain reaction (qRT-PCR) assays

Larvae were maintained in the 28.5°C incubator in petri dishes containing a volume of 45ml of fish water, with up to 60 larvae in each of four dishes. Where the larvae were to be treated with clonidine or DMSO, these drugs were added to the petri dish at 5 dpf. Where the larvae were to be treated with phenylephrine and isoproterenol, these drugs were added to the petri dish 50min prior to caffeine application. All drugs were applied to give the same final concentrations as in the sleep/wake assays. Caffeine/PTZ or water vehicle were applied at 6 dpf. After 1hr of caffeine/PTZ treatment, larvae were culled by addition of 8ml 25X tricaine (see Key Resources table) to each petri dish, and groups of ∼15-37 larvae were pipetted into 1.5ml Eppendorf tubes. Excess fish water was removed with a fine-tipped plastic pastette and sample tubes were frozen in isopentane on dry ice. Samples were then transferred to a −80°C freezer.

### qRT-PCR for measurement of *c-fos* mRNA levels

RNA isolation was performed on larval samples by homogenisation in TRIzol and treatment with chloroform. After centrifugation at 12,000g, the aqueous phase (containing RNA) was treated with 2-propanol and re-centrifuged at 12,000g. The RNA pellet was washed with 75% ethanol and resuspended in nuclease-free water. RNA quality was checked using Nanodrop. Only samples with a 260/280nm ratio of at least 1.8 (indicating minimal protein contamination) and a 260/230nm ratio of at least 1.9 (minimal phenol contamination) were used for analysis.

AffinityScript Reverse Transcriptase was used for reverse transcription of RNA. For each resulting sample of complementary DNA, levels of *fosab* (*c-fos)* were measured for three aliquots and of the housekeeping gene *ef1α* for another three aliquots, using GoTaq qPCR Master Mix, in a CFX96 Real-Time System BioRad Thermal Cycler. In zebrafish there are two paralogues to mammalian *c-Fos*: *fosaa* and *fosab*. The protein Fosab is the less divergent, with more highly conserved key regulatory phosphorylation sites (Kubra et al., 2022). The primers used for amplification of *fosab* (*c-fos)* and *ef1α* were per Reichert et al. (2019). The “quantification cycle” of *c-fos* from each sample was measured as the number of PCR cycles taken to reach the threshold level of fluorescence detection. This was then normalised to the quantification cycle of *ef1α* for the sample, giving the “delta quantification cycle” measure, “dCt”. The *c-fos* dCt of each sample was then normalised to the dCt measure of control sample(s), to give the “delta dCt” measure, “ddCt”. The relative *c-fos* expression for each sample versus control was then calculated as 2^-ddCt^.

### F0 KO zebrafish

Filial generation zero (F0) *dopamine β-hydroxylase* (*dbh*) knockout (KO) larvae were generated using a CRISPR/Cas9 F0 KO injection method (Kroll et al., 2021). To construct each guide RNA, 1µl of 200µM CRISPR RNA (crRNA) was annealed with 1µl of 200µM trans-activating CRISPR RNA (tracrRNA), in a mixture with 1.28µl of duplex buffer, at 95°C for 5min (see Key Resources table). 1µl of each guide RNA was then separately assembled with 1µl Cas9 nuclease at 37°C for 5 min to create a ribonucleoprotein complex. Eggs were injected at the 1-cell stage, shortly after laying, with ∼1nl of a mixture of three different ribonucleoprotein complexes. The three guide RNAs targeted different exons in the *dbh* gene to give a high chance of mutagenesis. The guide RNA target sequences were as follows: sequence 1: 5’-GACGCTGGTTTGCCTATGGG- 3’ (within exon 5), sequence 2: 3’-CGGGGGGGAATGGCCATCAC-5’ (within exon 6), and sequence 3: 3’- GGGACGGGGTGTCTGGACGC-5’ (within exon 3). Exons 5 and 6 were targeted because they are asymmetric (i.e., their base pair length is not a multiple of 3), increasing the likelihood of frameshift mutations in cases of exon skipping. Exon 3 was targeted because a mutation within this exon can give rise to non-functional Dbh (Singh et al., 2015).

Control eggs were injected with Cas9 assembled with non-targeting guide RNAs whose sequences were not predicted to match any genomic locus (see Key Resources table). Injected embryos were reared at 28.5°C.

### Deep sequencing of the *dbh* gene in F0 KO larvae

Illumina Miseq was used to estimate the rate of successful mutation of *dbh* copies in the F0 KOs, using MiSeq Reagent Nano Kit v2 (300 Cycles) (MS-103–1001), as per Kroll et al. (2021). Of the 29 *dbh* F0 KO larvae used to characterise the *dbh* F0 KO sleep/wake phenotype (Fig. S6), ten were selected for sequencing. Two control-injected larvae were also selected. Selection was made before inspection of behavioural data. Selected larvae were culled by tricaine overdose and pipetted into individual PCR tubes, from which fish water was then removed using a fine-tipped pastette. The PCR tubes were then frozen at −20°C. DNA extraction was performed on the 12 individual larvae using the HotSHOT method: 50µl of 1x base solution (see Key Resources table) was added to each larva before incubation for 30min at 95°C, then, after cooling, 50µl of 1x neutralisation solution (see Key Resources table) was added to each tube. The resulting DNA samples were diluted 2.5x with ddH_2_O and stored at −20°C for subsequent PCR.

PCR amplification was conducted for each of the three CRISPR-targeted regions for each DNA sample. Each PCR well contained: 1.20µl DNA template, 8.86µl nuclease-free water, 3.00µl Phusion High-Fidelity Reaction Buffer, 0.3µl 10mM deoxynucleoside triphosphates (dNTPs), 0.75µl 10µM forward primer, 0.75µl 10µM reverse primer, and 0.15µl Phusion High-Fidelity DNA Polymerase (see Key Resources table for all sources). The PCR program used was 95°C for 5min followed by 40 cycles of: 95°C for 30s, 60°C for 30s and 72°C for 30s, then 72°C for 5min and 10°C until collection. The following three pairs of forward and reverse primers were used, for sequences 1, 2 and 3, respectively. The Miseq adaptor arm sequence is shown, followed by the *dbh*-specific sequence (underlined):

5’-TCGTCGGCAGCGTCAGATGTGTATAAGAGACAG<u>ACTGTCATGGAACTACAGGGCT</u>-3’

5’-GTCTCGTGGGCTCGGAGATGTGTATAAGAGACAG<u>AAGGAGAGGGTTGTGGTAATGA</u>-3’

5’-TCGTCGGCAGCGTCAGATGTGTATAAGAGACAG<u>GGGCATTCGTTTATGGTACAGT</u>-3’

5’-GTCTCGTGGGCTCGGAGATGTGTATAAGAGACAG<u>TGGCTTGAGTGAAGTGCAGTAT</u>-3’

5’-TCGTCGGCAGCGTCAGATGTGTATAAGAGACAG<u>GCTCAATATATCCCGTCTCCAG</u>-3’

5’-GTCTCGTGGGCTCGGAGATGTGTATAAGAGACA<u>GTTATTTGTAATGTGCGAGTGGC</u>-3’

PCR product length was verified on a selection of three PCR products (and one control containing no PCR product) for each set of primers. Gel electrophoresis was performed using UltraPure Agarose and GelRed, with a 100bp DNA ladder and xylene cyanol loading dye. PCR product concentration was then measured for a selection of two PCR products for each set of primers using Qubit (dsDNA Broad Range Assay) and diluted as needed with ddH_2_O to a final DNA concentration of 15-25ng/µl. ExoSap-IT cleanup was then performed on all samples to degrade remaining primers and nucleotides.

Sequencing data was analysed per Kroll et al., 2021. Reads from one of the scrambled-injected controls were used to normalise mutation counts, so that misalignments present in the control were not considered to be Cas9 mutations in the F0 KOs. The scrambled-injected control from column 12 of the PCR plate was used for normalisation, as the column 11 control appeared to have been contaminated with DNA from column 10.

Of the 46 *dbh* F0 KOs used to investigate the effect of clonidine on these larvae (Fig. 6), ten were randomly selected for sequencing. Two control-injected larvae were also randomly selected. Sequencing was performed as above (per Kroll et al., 2021), with the exception that BAM files were not filtered prior to the inputting of fastq files to ampliCan, as sense-checking using Integrative Genomics Viewer (IGV) indicated that valid reads were being excluded by the filtering process.

### Statistical analysis

#### Statistical analyses were performed in MATLAB

For sleep/wake assays where two variables were manipulated, rebound sleep was compared between conditions using two-way ANOVA (e.g., stimulant treatment x noradrenergic status). Where one variable was manipulated (e.g., noradrenergic status), one-way ANOVA was used.

Differences in qRT-PCR measurements of *c-fos* expression were statistically analysed across conditions using the Wilcoxon two-sample test, at the level of the dCt metric (Yuan et al., 2006). This nonparametric test was appropriate given the small sample sizes, making no assumption of data normality.

Linear regression analysis was performed to assess the relationship between *c-fos* expression and rebound sleep across drug conditions, with calculation of the R^2^ goodness-of-fit measure.

The Kolmogorov-Smirnov two-sample test was used to assess the difference between the frequency distributions of sleep/wake bout lengths of *dbh* F0 KOs and controls.

## Results

### Pre-treatment of larvae with clonidine facilitates drug-induced rebound sleep

To assess the effects of suppressing noradrenergic transmission during neuronal hyperactivation on subsequent homeostatic rebound sleep, we induced rebound sleep in larval zebrafish with acute stimulant exposure while also pharmacologically targeting α_2_-adrenoceptors (Fig. 1). α_2_-adrenoceptors are G-protein-coupled-receptors that principally bind G_i_-proteins to inhibit adenylyl cyclase activity (Perez, 2020; Jasper et al., 1998). As such, activation of α_2_-adrenoceptors tends to inhibit neuronal activity, including autoinhibiting the LC, causing sedation (Nelson et al., 2003). Indeed, clonidine has been shown to enhance sleep in zebrafish (Rihel et al., 2010). Accordingly, following clonidine administration at 5 days post fertilisation (dpf), and prior to exposure to stimulant drugs, sleep levels were increased (Fig. 1a, 1c). After a ∼20hr exposure to clonidine, larvae were then treated with either caffeine (Fig. 1a, 1b) or PTZ (Fig. 1c, 1d) for 1hr to acutely increase neuronal activity and generate rebound sleep upon wash-off. As expected, treatment with either caffeine (Fig. 1a) or PTZ (Fig. 1c) alone caused sleep levels to be greatly increased during the rebound period from the end of the drug wash-off to lights off at ZT14.

This rebound sleep is thought to reflect the greater sleep need caused by enhanced neuronal activity during stimulant exposure (Reichert et al., 2019).

In both experiments, there was also a main effect of prior clonidine treatment on boosting subsequent sleep in the rebound period. This effect trended towards statistical significance in the caffeine protocol (p=0.056, Fig. 1b) and was statistically significant (p<0.05) for the PTZ experiment (Fig. 1d). One explanation for this could be that clonidine washed out of the larval brain less quickly than caffeine/PTZ, continuing to agonise α_2_-adrenoceptors somewhat into the rebound period. However, inspection of clonidine-treated larvae that were not given a stimulant drug (blue traces in Fig. 1a, 1c) reveals that their daytime sleep levels were only heightened versus controls (gray traces) when clonidine was present in the fish water. Directly after wash-off, sleep of clonidine-only treated animals was similar to control levels, suggesting successful rapid wash-off. To confirm the rebound sleep effects of clonidine in caffeine-treated larvae, the experiment was simplified and repeated with only two experimental conditions: 96 larvae were treated at 5 dpf with either clonidine or DMSO vehicle and then exposed to caffeine for 1hr on the following morning at 6 dpf (Fig. S1). Larvae treated with clonidine showed significantly higher levels of rebound sleep following caffeine wash-off than DMSO-treated larvae (p=8.8 x 10^-8^, F(1,94) = 33.67, one-way ANOVA). These results not only demonstrate that noradrenergic arousal is not required for neuronal activity-dependent rebound sleep but also suggest that reduced noradrenergic tone may in fact enhance rebound sleep.

### *c-fos* induction by neuronal activity-promoting drugs is greater following pre-treatment with clonidine

In zebrafish, both PTZ-and caffeine-induced rebound sleep are positively correlated with the neuronal activity driven during stimulant exposure (Reichert et al., 2019). Clonidine is a sedative and was predicted to dampen neuronal activity during stimulant exposure, yet it enhanced rebound sleep. Therefore, we next investigated the effects of clonidine on stimulant-induced neuronal activity by assessing expression of the immediate early gene *c-fos*. Brain-wide *c-fos* expression is enhanced upon waking and after stimulation (Cirelli and Tononi, 2000) and is a widely-used indicator of neuronal activity, including in zebrafish. In control experiments, caffeine-treated larvae showed on average 71-fold higher *c-fos* expression than water-treated larvae (Fig. 2a, S2a), consistent with previous observations (Reichert et al., 2019). However, contrary to expectations, when larvae were co-treated with caffeine and clonidine, *c-fos* expression was elevated even further, being 47% higher than in larvae treated only with caffeine (Fig. 2b, S2b). Notably, there was a strong correlation (R^2^ = 0.985) between the relative *c-fos* expression induced by combinations of clonidine and caffeine and the associated rebound sleep (Fig. 2c), consistent with previous findings in zebrafish that rebound sleep duration correlates with *c-fos* levels induced during drug exposure (Reichert et al., 2019).

**Fig. 2.**
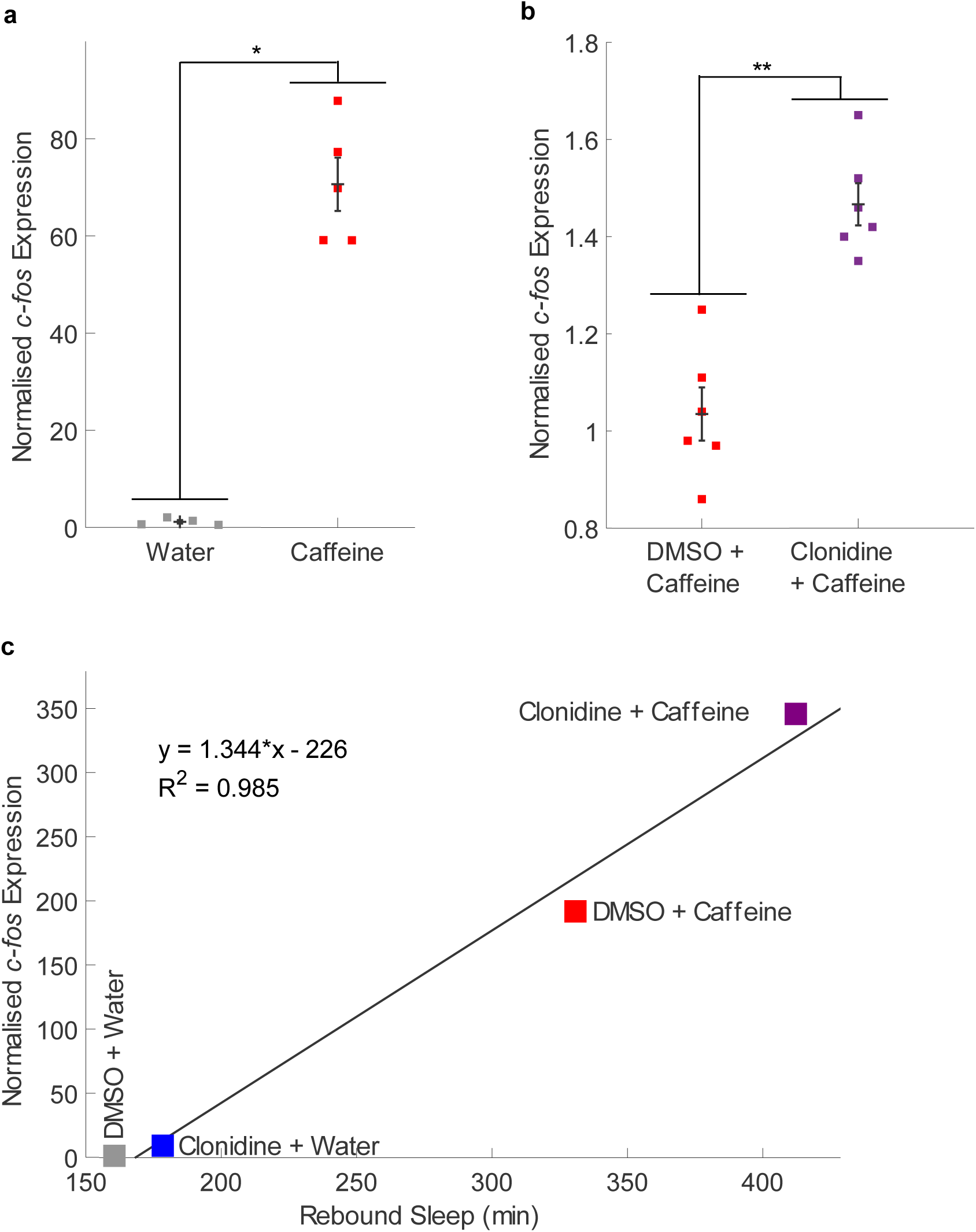
*c-fos* expression is higher in larvae following combined treatment with clonidine and caffeine than following caffeine alone. **a** qRT-PCR on groups of ∼20 larvae (n = 4 and n = 5 biological replicates per condition) reveals that larvae treated with caffeine had a significant, 71-fold increase in *c-fos* expression compared to water-treated larvae (*p<0.05, two tailed Wilcoxon rank sum test performed on the “dCt” metric, see Fig. S2a). **b** *c-fos* expression of larvae soaked in clonidine before and during caffeine exposure was significantly higher by 47% than in larvae exposed to caffeine alone (n = 6 biological replicates per condition, *p<0.01, see Fig. S2b). **c** The relative *c-fos* expression induced by different combinations of vehicle, clonidine and caffeine is positively, linearly correlated (R^2^ = 0.985) with the total rebound sleep induced by these drugs (see Fig. 1a-b). qRT-PCR was performed on groups of 37 larvae (see Fig. S2c). Each square in **a-c** is the mean of three technical replicates

To test whether clonidine’s enhancement of caffeine-induced *c-fos* expression was drug-specific, we also measured *c-fos* expression in larvae following treatment with clonidine and PTZ. As observed for caffeine, treatment with clonidine and PTZ further enhanced *c-fos* expression compared to PTZ treatment alone (Fig. 3a, S3). As in the clonidine/caffeine experiments, there was a strong correlation (R^2^ = 0.993) between the relative *c-fos* expression levels in the different clonidine/PTZ treatment conditions and their associated amount of rebound sleep (Fig. 3b). Thus, depressing the noradrenergic system by activating α_2_-adrenoceptors actually enhances the expression of *c-fos*, and the level of *c-fos* induction predicts the duration of subsequent rebound sleep.

**Fig. 3.**
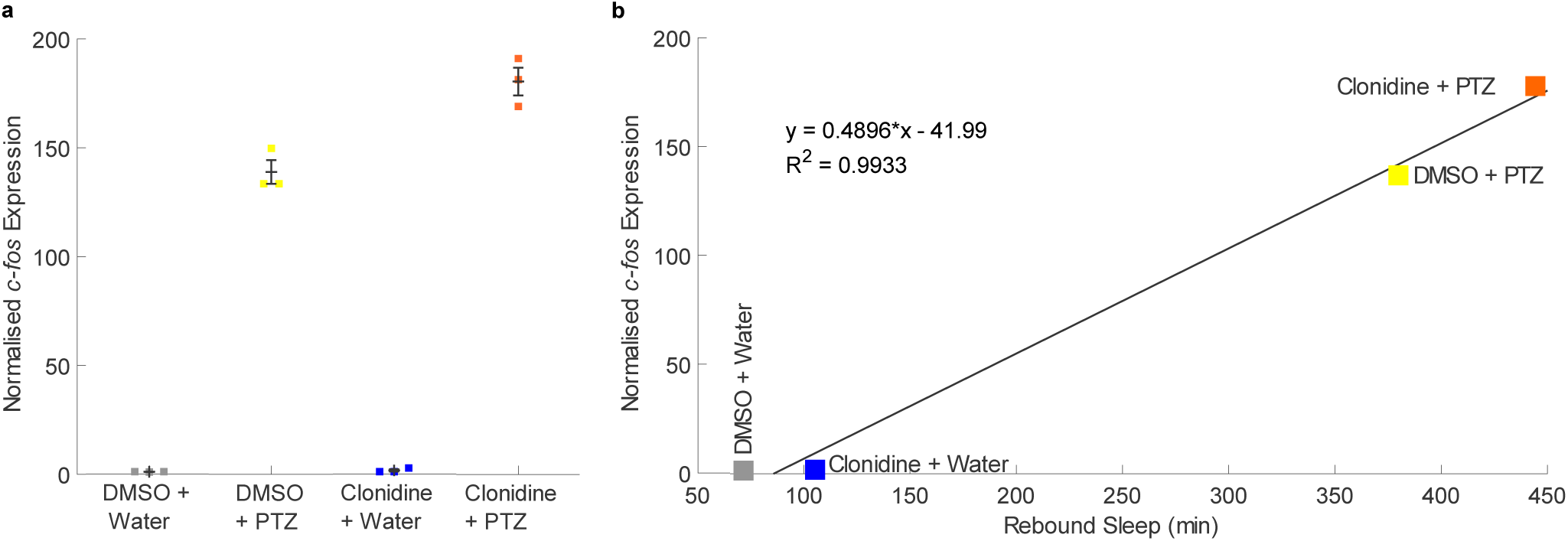
Rebound sleep levels correlate with *c-fos* expression across different clonidine/PTZ treatment combinations. **a** qRT-PCR on groups of ∼17 larvae (n=3 biological replicates per condition) reveals that larvae treated with both clonidine and PTZ had a trend towards higher *c-fos* expression than those treated with PTZ alone (see also Fig. S3a). **b** The mean *c-fos* expression induced by each drug combination is strongly positively correlated (R^2^ = 0.993) with the amount of rebound sleep induced by each drug condition (see Fig. 1c-d). Each square in **a** is the mean of three technical replicates

### Stimulation of α1-and β-adrenoceptors with isoproterenol and phenylephrine does not boost neuronal activity-induced rebound sleep

Since the inhibition of noradrenergic signalling with clonidine enhanced stimulant-induced *c-fos* expression and rebound sleep, we next tested the effects of activating noradrenergic transmission by agonising both α_1_-and β-adrenoceptors while inducing rebound sleep with caffeine exposure. Phenylephrine is an agonist of the principally G_q_-coupled α_1_-adrenoceptors (Perez, 2020) and thus tends to enhance neuronal excitability. Isoproterenol is an agonist of β-adrenoceptors, which couple to G_s_-proteins to enhance neuronal activity via the stimulation of adenylyl cyclase (Perez 2020), and has been shown to reduce sleep in zebrafish (Rihel et al., 2010). Larvae (6 dpf) were pre-treated at ZT0 with a cocktail of phenylephrine and isoproterenol to activate both α_1_- and β-adrenoceptors, followed by a 1hr caffeine exposure at ZT1 and then wash-off of all drugs (Fig. 4a). Although caffeine significantly induced rebound sleep, the addition of isoproterenol and phenylephrine did not enhance rebound sleep (Fig. 4a-b). In fact, groups pre-treated with isoproterenol and phenylephrine showed marginally lower rebound sleep levels than water-treated groups in both caffeine and control conditions (Fig. 4b), though the effect was not statistically significant (p = 0.21).

**Fig. 4.**
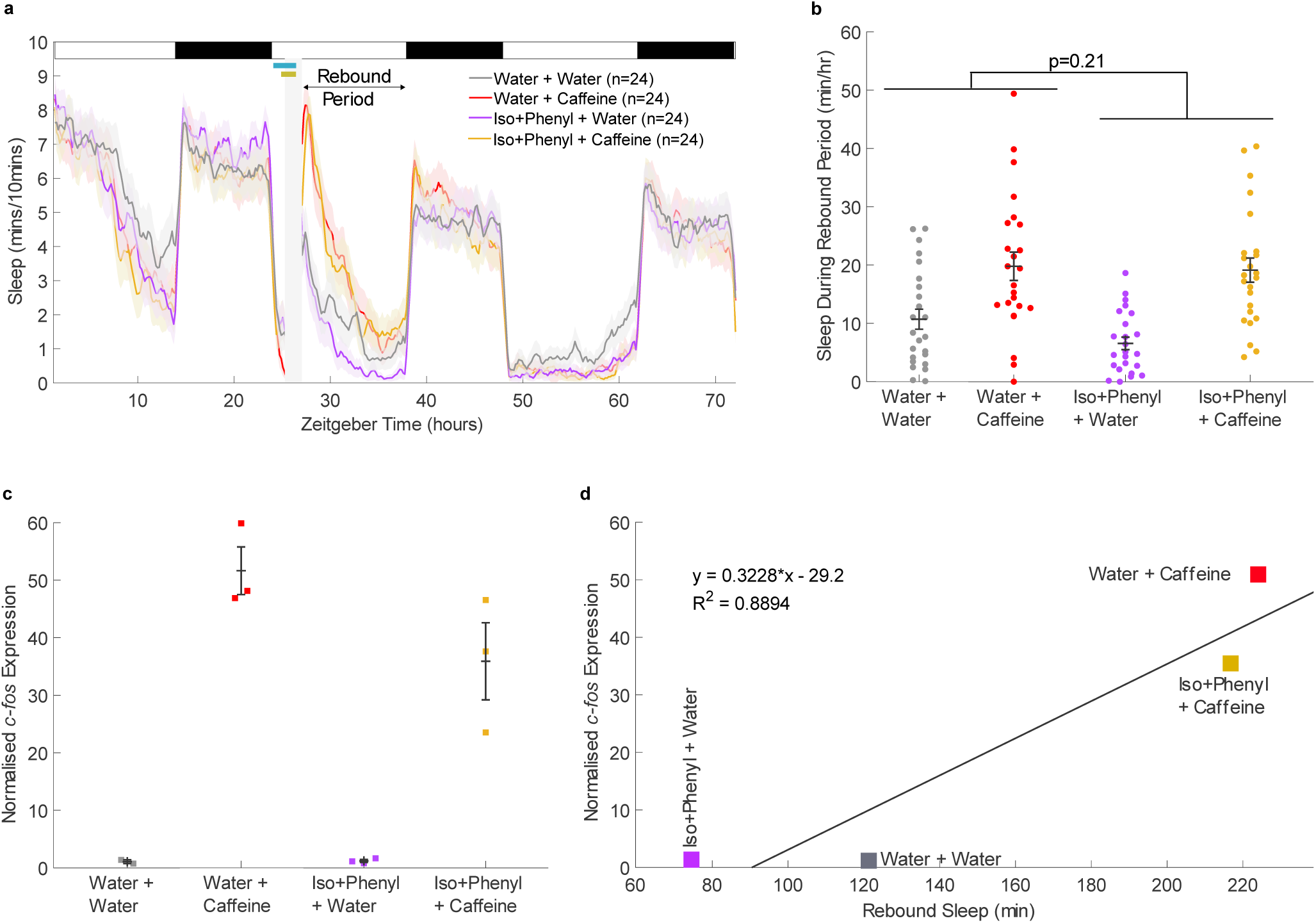
Activating noradrenergic transmission with isoproterenol and phenylephrine marginally depresses caffeine-induced *c-fos* expression. **a** Sleep traces for larvae exposed to combinations of 10µM isoproterenol + 10µM phenylephrine (“Iso+Phenyl”) and/or caffeine. At top left, the pale blue horizontal bar shows the isoproterenol+phenylephrine exposure window while the gold bar indicates the presence of caffeine. The post-drug rebound sleep period of **a** is summarised for each larva in **b**. Caffeine treatment significantly increased subsequent rebound sleep (p=1.2 x 10^-7^, F(1,92) = 32.89), while the effect of isoproterenol+phenylephrine treatment was not significant (p=0.210, F (1,92) = 1.6). There was no significant interaction between isoproterenol+phenylephrine and caffeine treatment (p=0.358, F(1,92) = 0.85), based on a two-way ANOVA (caffeine treatment x isoproterenol+phenylephrine treatment). **c** qRT-PCR on groups of ∼18 larvae reveals that each group of larvae pre-treated with isoproterenol+phenylephrine and then caffeine (n=3 biological replicates) had lower relative *c-fos* expression than the groups of larvae treated with water and then caffeine (n=3 biological replicates); see also Fig. S4a. Each square is the mean of three technical replicates. **d** The average relative *c-fos* expression induced by each condition is strongly positively correlated (R^2^ = 0.889) with the total rebound sleep that was induced by the same drug condition (from **a-b**)

We then tested the effects of isoproterenol and phenylephrine treatment during caffeine exposure on the induction of *c-fos* expression. In contrast to the enhancement of *c-fos* expression observed when noradrenergic tone was dampened with clonidine, caffeine-induced *c-fos* expression was lower in groups pre-treated with isoproterenol and phenylephrine than in water-treated controls (Fig. 4c). We repeated this *c-fos* measurement with six additional groups of larvae treated with isoproterenol and phenylephrine and six groups treated with water and confirmed that caffeine-induced *c-fos* expression was on average collectively lower among groups pre-treated with isoproterenol and phenylephrine, but the effect only trended toward significance (p = 0.077, Fig. S4b). However, as in the clonidine experiments, there was a strong positive correlation (R^2^ = 0.889) between the relative *c-fos* expression induced by the different drug treatments and the duration of rebound sleep (Fig. 4d), again suggesting a relationship between the magnitude of *c-fos* induction during stimulant treatment and the sleep pressure generated.

### *dbh* F0 KO larvae phenocopy the high sleep levels of *dbh^-/-^* mutants

To complement our pharmacological manipulations of the noradrenergic system and ensure that the effects we had observed were not drug-specific (e.g., off-target effects), we used a genetic knock-out approach to disrupt the *dopamine β-hydroxylase* (*dbh*) gene, which is necessary for noradrenalin synthesis. To eliminate *dbh* function, we injected zebrafish eggs with Cas9 nuclease assembled with guide RNAs that targeted three loci within the *dbh* gene (see Methods; Kroll et al., 2021). The resulting *dbh* F0 KO larvae were used for experiments at 5-8 dpf.

To verify that *dbh* function was successfully disrupted in most if not all cells of the F0 KO larvae, we performed deep-sequencing on larval samples and ascertained the frameshift and mutation rates for each of the three targeted loci within the *dbh* gene. For 10 sequenced F0 KOs (taken at the end of the experiment, see Fig. S6), the proportion of reads that harboured either mutations or frameshift mutations exceeded 50% at each locus in most larvae. One larva, F0 KO 9, was an exception with no mutated reads at any locus, likely due to experimenter error (e.g., an uninjected egg that was trapped in the transfer pipette) (Fig. S5a). Considering all three targeted loci together, 9/10 of the F0 KO larvae had at least 50% frameshifted copies of *dbh*, and 7/10 had above 80% (Fig. S5b). This high rate of success, which does not take into account the likelihood that non-frameshifting mutations are also deleterious, indicates that most F0 KO larvae were largely, if not completely, functionally null for *dbh* in most or all cells.

Previous studies have shown that *dbh* knockout zebrafish (*dbh^-/-^*) have elevated baseline sleep, especially during the day (Singh et al., 2015). This reflects the inability of *dbh^-/-^* mutants to synthesise the arousal-promoting neurotransmitters noradrenalin and adrenalin (which is synthesised from noradrenalin). We hypothesised that if our *dbh* F0 KOs were loss-of-function, they would similarly show enhanced sleep, particularly during the day when the arousal systems of diurnal species are most active. Tracking *dbh* F0 KOs from 5 dpf over several day/night cycles revealed that they had significantly elevated sleep levels, especially during the day, with *dbh* F0 KOs sleeping on average 50% of the time at 6 dpf (Fig. S6a-b), versus 15% for controls. *dbh* F0 KOs were unable to sustain wakefulness for long periods, showing significantly shorter wake bouts and a trend towards longer sleep bouts than controls (Fig. S6c-d).

To ascertain more carefully how closely *dbh* F0 KOs recapitulated the sleep phenotype of published *dbh^-/-^* null mutants, we compared the sleep parameters of *dbh* F0 KOs to those of stable *dbh^-/-^* knockout animals as reported in Singh et al. (2015) (underlying data courtesy of David Prober). On average, 6 dpf *dbh* F0 KOs showed +233% higher total daytime sleep compared to control larvae, similar to the +225% elevation of daytime sleep in *dbh^-/-^* null mutants (Fig. S7a). Similar results were found in night-time sleep, with *dbh* F0 KO larvae having an average +49% increase in total night-time sleep (compared to +58% in *dbh^-/-^* null mutants) (Fig. S7b). As in *dbh^-/-^* null mutants, the day and night increases in sleep were due to both an increase in the number and length of sleep bouts. In the day, *dbh* F0 KO larvae had an increase in sleep bout number (+107%, compared to +201% in *dbh^-/-^*mutants) and sleep bout length (+63%, compared to +17% in *dbh^-/-^*mutants) (Fig. S7c, S7e). This discrepancy in daytime effect sizes could reflect the different lighting and temperature conditions in which the larvae were raised (in two different labs on separate continents) as well as the potentially incomplete knockout of *dbh* in F0 KOs. At night, *dbh* F0 KO larvae and *dbh^-/-^* mutants showed broadly similar elevations of sleep bout number (+17% and +27% respectively) and sleep bout length (+26% and +30%) (Fig. S7d, S7f), demonstrating a high degree of similarity in sleep phenotypes between *dbh* F0 KOs and *dbh^-/-^* mutants at night.

Taken together, the sequencing data combined with the similarity between *dbh* F0 KO and stable *dbh^-/-^*knockout animals’ sleep phenotypes suggests that *dbh* F0 KOs lack Dbh function and are therefore, like *dbh^-/-^* mutants (Singh et al., 2015), depleted of noradrenalin.

### *dbh* F0 KOs show enhanced caffeine-induced *c-fos* expression and robust rebound sleep

Having verified that our CRISPR/Cas9 technique was generating effective *dbh* knockouts, we used *dbh* F0 KOs in an assay of caffeine-induced rebound sleep to test the effect of genetic noradrenergic impairment. An important distinction in this experiment versus our pharmacological noradrenergic manipulations is that the genetic noradrenergic impairment is persistent, whereas pharmacological activation of adrenoceptors should cease after drug wash-off. As such, here we observed the ongoing effects of noradrenergic impairment on rebound sleep, rather than the after-effects. Based on the effects of pharmacological manipulation of adrenoceptors, we predicted that rebound sleep would occur robustly in *dbh* F0 KOs. Indeed, after caffeine wash-off, *dbh* F0 KOs showed an average increase of 186min (+58%) of rebound sleep versus water-treated *dbh* F0 KOs (Fig. 5a), indicating that drug-induced rebound sleep can still occur without noradrenalin (Fig. 5a-b).

**Fig. 5.**
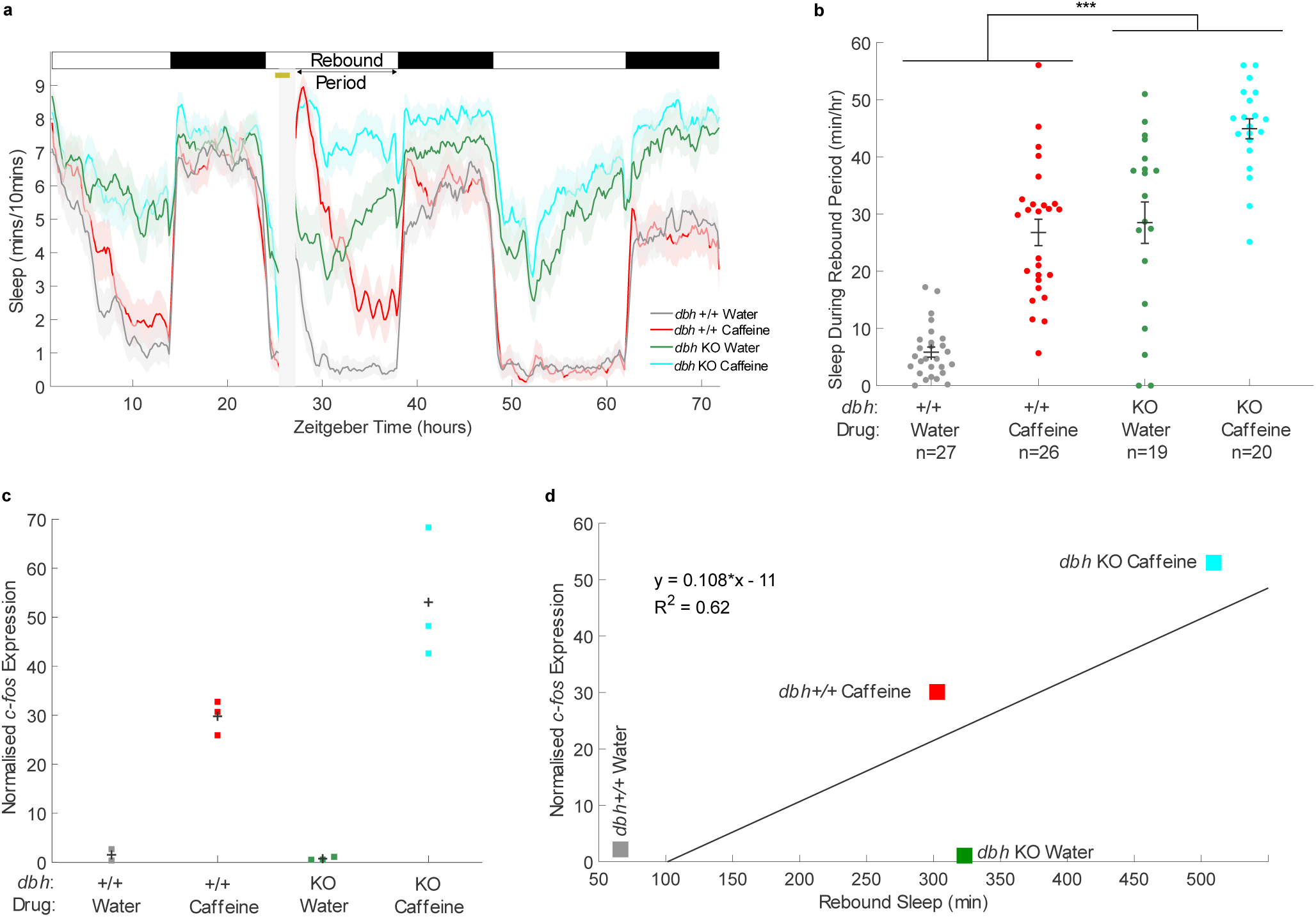
Caffeine triggers *c-fos* expression more strongly in *dbh* F0 KOs. **a** Sleep traces for *dbh* F0 KO and control-injected larvae exposed to either caffeine or water. The post-drug rebound sleep period of **a** is summarised for each larva in **b**. Caffeine treatment significantly increased subsequent rebound sleep (p=5.3 x 10^-13^, F(1,88) = 71.54), as did *dbh* KO (p=1.2 x 10^-14^, F(1,88) = 85.59), but there was no significant interaction between *dbh* genotype and caffeine treatment (p=0.31, F(1,88) = 1.04), based on a two-way ANOVA (caffeine treatment x *dbh* genotype). **c** qRT-PCR on groups of ∼15 larvae revealed that each group of *dbh* F0 KO larvae treated with caffeine (n=3 biological replicates) showed greater relative *c-fos* expression than the groups of control larvae treated with caffeine (n=3 biological replicates); see also Fig. S8a. Each square is the mean of triplicate technical replicates. **d** There is a trend towards a positive correlation between *c-fos* expression and subsequent rebound sleep levels (R^2^ = 0.62), but water-treated *dbh* F0 KO larvae do not conform to this trend, showing high sleep levels despite low *c-fos* expression. ***p<0.001

Furthermore, there was not a significant interaction effect between genotype and caffeine treatment on rebound sleep (p=0.31, caffeine treatment x *dbh* genotype interaction, two-way ANOVA), again suggesting that stimulant-induced rebound sleep can occur independently of *dbh*.

To assess how the loss of *dbh* impacted neuronal activity during the stimulant treatment, *c-fos* expression immediately following caffeine treatment was measured in both *dbh* F0 KOs and controls. As for larvae with pharmacologically compromised noradrenergic systems (via activation of α_2_-adrenoceptors with clonidine), *c-fos* expression was elevated in *dbh* F0 KOs treated with caffeine versus caffeine-treated wild-type controls (Fig. 5c). However, unlike in the clonidine experiments (Fig. 2c, 3b), there was only a weak correlation between *c-fos* expression and sleep across all *dbh* conditions (Fig. 5d). This difference from the pharmacological experiments is likely due to the high sleep levels during the rebound phase of *dbh* F0 KOs that were exposed only to water, despite the low induction of *c-fos* expression in these animals. This indicates that, unsurprisingly, high *c-fos* expression during the stimulant window is not a prerequisite for the high levels of baseline sleep seen in *dbh* F0 KOs. Nonetheless, exposure to caffeine does induce *c-fos* expression and subsequent rebound sleep in animals that lack noradrenalin.

### Clonidine’s sedative effects are not mediated solely by α_2_-autoreceptor suppression of noradrenalin release

One model for how the α_2_-adrenoceptor agonist dexmedetomidine initiates sedation is by primarily activating auto-inhibitory α_2_-adrenoceptors found presynaptically on LC neurons, thereby suppressing release of noradrenalin (Nelson et al., 2003). However, other work indicates that α_2_-adrenoceptors can act as heteroreceptors, sitting presynaptically on non-noradrenergic neurons to inhibit release of glutamate (Harris et al., 2018; Shields et al., 2009). Additionally, α_2_-adrenoceptors can sit post-synaptically and even be excitatory (Harris et al., 2018; Jasper et al., 1998). Indeed, Hu et al. (2012) found that *Dbh^-/-^*mice are hypersensitive to dexmedetomidine, indicating that the sedative effects of this α_2_-adrenoceptor agonist do not rely solely on the inhibition of noradrenergic release. We reasoned that if clonidine causes sedation primarily via suppression of noradrenalin release, then the sedative effects of clonidine should be blunted in *dbh* F0 KO larvae. Alternatively, if clonidine enhances sleep independently of its inhibition of noradrenergic release, the sleep-inducing effect of clonidine should occur additively, on top of the elevated baseline sleep phenotype seen in *dbh* F0 KOs.

Applying clonidine to 5 dpf larvae caused daytime sleep levels to rise substantially in both *dbh* F0 KO and control-injected larvae (Fig. 6a-b), with a significantly boosted sleep level in *dbh* F0 KOs (Fig. 6b, p=0.0012, one-way ANOVA). Thus, clonidine’s sedative effects are not solely due to the suppression of noradrenalin release, as additional sedation was induced in *dbh* knockout animals that lack noradrenalin. There was a significantly stronger effect of clonidine in control-injected larvae (p=0.0034, *dbh* genotype x clonidine treatment interaction, two-way ANOVA), which is consistent with clonidine’s sedative effects being at least partially mediated by suppression of the noradrenergic system; however, baseline daytime sleep levels are already very elevated in *dbh* F0 KOs, capping the sedative effect that could be achieved by the addition of clonidine, and so limiting interpretation.

**Fig. 6.**
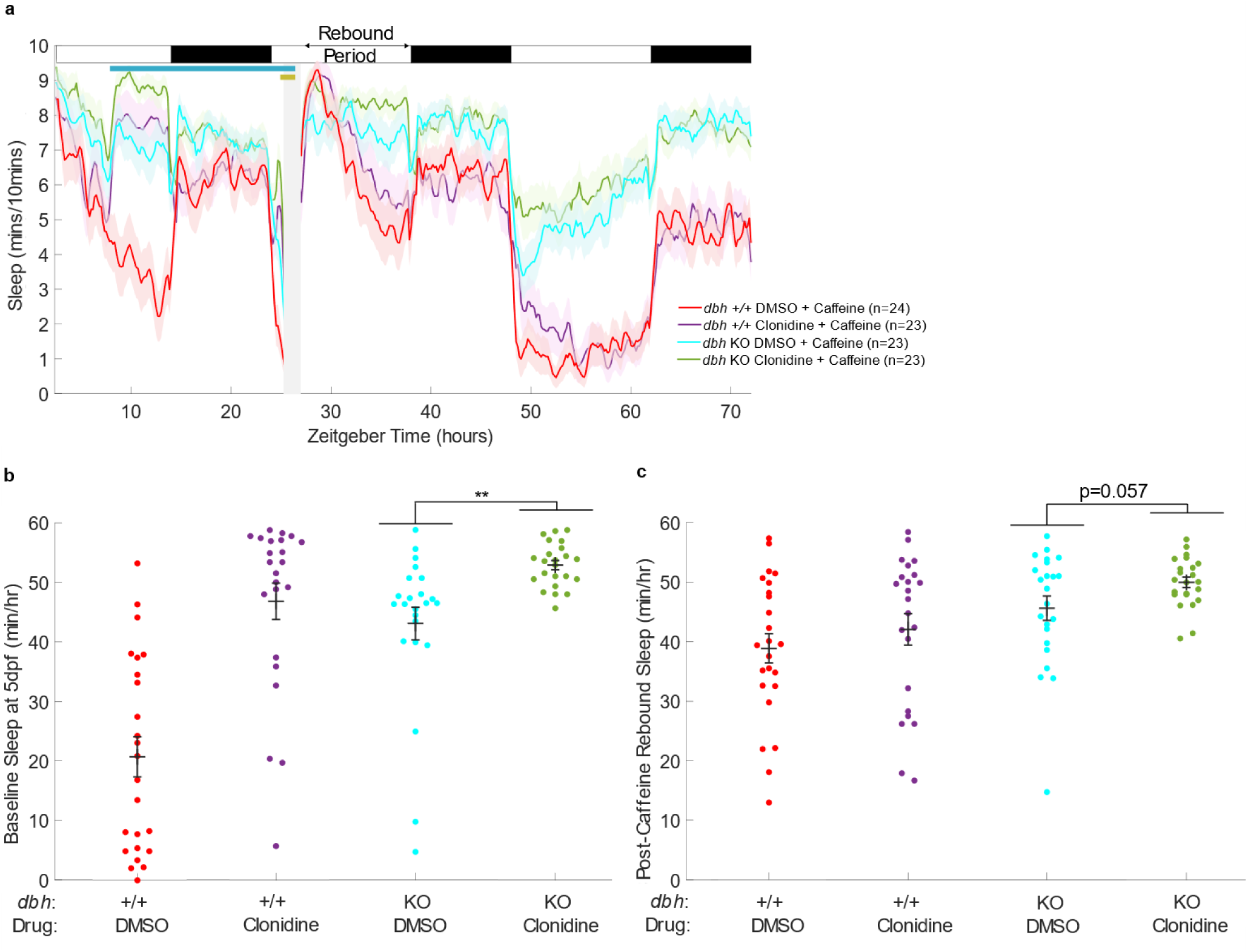
Clonidine enhances sleep, and rebound sleep, in *dbh* F0 KO larvae. **a** Sleep traces for *dbh* F0 KO and control-injected larvae exposed to clonidine/DMSO and caffeine. **b** At 5 dpf from clonidine treatment until lights-out, clonidine significantly boosted sleep levels (p=2.3 x 10^-9^, F(1,89) = 44.25), as did loss of *dbh* (p=9.9 x 10^-7^, F(1,89) = 27.65). There was also a significant interaction between clonidine treatment and *dbh* genotype (p=0.0034, F(1,89) = 9.06), based on a two-way ANOVA (*dbh* genotype x clonidine treatment). Among *dbh* F0 KOs larvae, clonidine treatment significantly boosted sleep (p=0.0012, F(1,44) = 11.93), based on a one-way ANOVA. The post-caffeine rebound sleep period is summarised for each larva in **c**. *dbh* F0 KOs had a significant increase of sleep (p= 0.0009, F(1,89) = 11.86), while the effect of clonidine treatment trended to increase sleep (p=0.079, F(1,89) = 3.15). There was no significant interaction between *dbh* genotype and clonidine treatment (p=0.786, F(1,89) = 0.07), based on a two-way ANOVA (*dbh* genotype x clonidine treatment). Among *dbh* F0 KO larvae, clonidine treatment boosted rebound sleep (p=0.0567, F(1,44) = 3.83), based on a one-way ANOVA. **p<0.01. Deep sequencing was used to verify the successful loss-of-function targeting of *dbh* in 10 randomly selected *dbh* F0 KO larvae: all animals had >93% (mean, 96%) of their amplified *dbh* copies frameshifted (see Fig. S9)

Following the pre-treatment of larvae with clonidine, we also induced homeostatic rebound sleep with acute exposure to caffeine, to test the effects of clonidine on subsequent rebound sleep in *dbh* F0 KOs. As in wild-type larvae, clonidine enhanced rebound sleep in *dbh* F0 KOs (Fig. 6c), with a trend towards significance (p=0.0567, one-way ANOVA). Indeed, clonidine’s rebound sleep-enhancing effects in *dbh* F0 KOs were not different from its effects in wild-type larvae (p=0.786, *dbh* genotype x clonidine treatment, two-way ANOVA), indicating that clonidine’s enhancement of rebound sleep may not arise from the after-effects of its α_2_-autoreceptor-mediated suppression of noradrenergic release.

## Discussion

Noradrenergic tone is highest during waking and promotes neuronal activity and behavioural arousal in vertebrate species including rodents and zebrafish (Carter et al., 2010; Wang et al., 2022). We therefore tested the effects of altering noradrenergic signalling in zebrafish on stimulant-drug-induced rebound sleep, which is hypothesised to be dependent on heightened neuronal activity (Reichert et al., 2019). Unexpectedly, pharmacological inhibition of noradrenergic signalling enhanced stimulant-induced homeostatic rebound sleep, while stimulant-induced *c-fos* expression was strongest in noradrenergic-compromised larvae. This enhancement of immediate early gene expression may thus underlie the increase in rebound sleep, for example by strengthening a sleep pressure signal, either on a brain-wide basis or in a key cell population. Alternatively, diminished noradrenergic arousal may de-potentiate widespread neuronal transmission, causing lingering quiescence into the rebound phase.

### Noradrenergic tone inversely modulates stimulant-induced *c-fos* expression

We found that stimulation of noradrenergic α_1_-and β-adrenoceptors with a cocktail of phenylephrine and isoproterenol slightly reduced *c-fos* induction by caffeine in zebrafish larvae and did not enhance subsequent rebound sleep. On the other hand, treatment of larvae with the α_2_-adrenoceptor agonist clonidine, a sedative, boosted *c-fos* induction by caffeine and enhanced rebound sleep. Likewise, *dbh* F0 KOs, which lack noradrenalin, showed elevated stimulant-induced *c-fos* expression and robust rebound sleep. These effects on the induction of *c-fos* are consistent with studies that identify *c-fos* expression as a measure of increases, as opposed to absolute levels, of neuronal activity. Indeed, c-Fos can show a refractory period after seizure induction, during which further seizures do not bring on c-Fos expression (Barros et al., 2015), and immediate early genes are not continually expressed in neurons that are chronically active (Hudson 2018). Rather, *c-fos* expression occurs in response to a change in stimulation, after which there may be self-inhibition of the *c-fos* promoter (Hudson, 2018). Such self-inhibitory regulation of *c-fos* expression could explain why *c-fos* induction is stronger when a stimulus is applied to an animal after a period of sensory deprivation (Cirelli and Tononi, 2000). During waking, because noradrenalin enhances the excitability of thalamic relay projections in mammals (Jones, 1991; Szymusiak and McGinty, 2008), a noradrenergic-compromised animal may be less aware of stimuli, akin to being sensorily deprived. As such, absolute levels of neuronal activity may not be higher in noradrenergic-compromised larvae that in control larvae following stimulant treatment, but the induction of *c-fos* may be stronger due to a greater magnitude of increase in neuronal activity. This prediction could be tested using larval zebrafish whole-brain neuronal imaging with genetically encoded calcium indicators to measure the ongoing neuronal activity during and after stimulant treatment. Another factor that may contribute to enhanced *c-fos* expression in noradrenergic-compromised larvae is that we performed our stimulant drug assay during the day, when *dbh* F0 KO larvae are much more likely to be asleep than wild-type controls. Thus, daytime drug administration will have caused a higher proportion of noradrenergic-compromised animals to undergo sleep-to-wake transitions, potentially bringing on a larger *c-fos* induction. Regardless of the precise mechanistic underpinnings, in our experiments, both genetic and pharmacological inhibition of noradrenergic signalling led to enhanced stimulant-induced *c-fos* expression.

### Magnification of immediate early gene induction may enhance a sleep pressure signal

Consistent with the findings of Reichert et al. (2019) that levels of pharmacologically-induced rebound sleep correlate with *c-fos* levels, we found a strong correlation between *c-fos* expression and sleep across noradrenergic/stimulant drug treatment combinations. One explanation for this could be that c-Fos protein, a transcription factor, drives expression of a homeostatic sleep pressure signal (Cirelli et al., 1995). Greater *c-fos* expression in noradrenergic-compromised, stimulant-treated larvae would then drive a stronger sleep pressure signal, enhancing rebound sleep. To test whether elevated *c-fos* expression plays a role in driving heightened rebound sleep, behaviour could be assayed in transgenic zebrafish larvae with inducible extra copies of the *c-fos* gene, which under this model would heighten rebound sleep following stimulant treatment. Conversely, animals with knock-down of *c-fos* would be expected to show blunted rebound sleep. If *c-fos* manipulations do indeed alter rebound sleep, additional experiments that restrict the overexpression or knockdown to particular subsets of neurons could be used to dissect whether distinct neuronal populations have particular roles in mediating sleep homeostasis. In addition, expression levels of many other immediate early genes including *Bdnf* and *Egr1* have been shown to correlate with homeostatic sleep pressure in mice (Vassalli and Franken, 2017) and are acutely and strongly induced by arousing drugs in zebrafish (Sabine Reichert, unpublished observation). Furthermore, the protein product of another immediate early gene, *Npas4*, was recently shown to help repair neuronal activity-induced DNA double strand breaks (Pollina et al., 2023), and in zebrafish, the build-up of neuronal DNA damage during waking has been shown to increase sleep pressure (Zada et al., 2021). Although it is unknown whether induction of these and other immediate early genes changes in response to manipulation of the noradrenergic system, their possible roles in regulating drug-induced rebound sleep in zebrafish larvae should be explored.

Alternatively, the correlation of the level of *c-fos* induction with subsequent rebound sleep may reflect altered activity of CREB, which mediates *c-fos* transcription in response to various stimuli (Ahn et al., 1998). Recent work in mice has demonstrated that CREB, in conjunction with the histone deacetylase HDAC4, acts downstream of the kinase SIK3 to regulate sleep (Kim et al., 2022; Zhou et al., 2022). Heightened *c-fos* induction during waking may cause changes in CREB’s interaction with HDAC4 and altered transcription of their targets as a function of sleep need. Such a model could be tested by modulating SIK3, HDAC4, or other components of this pathway in zebrafish and observing how drug-induced rebound sleep is affected.

### Heightened noradrenergic tone is not required for stimulant-induced *c-fos* expression or sleep rebound

How drug-induced neuronal activation leads to heightened rebound sleep is unclear; however, the neuropeptide galanin plays a critical role in the response to sleep pressure signals in zebrafish, functioning as an output arm of a sleep homeostat (Reichert et al., 2019). In mammals, a “flip-flop” model of sleep regulation posits that mutual inhibition between wake-promoting neurons such as those of the LC and sleep-promoting GABAergic/galaninergic neurons of the POA enables rapid and absolute transitions between sleep and wake (Saper et al., 2010). *dbh* F0 KOs lack noradrenalin, so noradrenergic tone is already supressed regardless of the drug treatment they receive. We found that control larvae showed a greater increase in rebound sleep after caffeine treatment (+237 min) than *dbh* F0 KOs (+186min), especially just after wash-off (Fig. 5a). This suggests that suppression of noradrenergic release is one mechanism involved in driving rebound sleep, consistent with a flip-flop model. In this interpretation, noradrenergic output cannot be further supressed in the *dbh* F0 KOs, explaining their reduced increase in sleep early in the rebound period compared to the control larvae. However, across the entire rebound period, both *dbh* F0 KOs and controls had statistically similar sleep rebound responses to caffeine, suggesting that release of noradrenalin from the LC during stimulant drug exposure is not necessary for rebound sleep to subsequently ensue. Indeed, the fact that administering caffeine to *dbh* F0 KOs enhances their rebound sleep at all, which was similarly observed in clonidine-treated larvae, indicates that noradrenergic tone during waking is not required for the generation of robust neuronal activity-induced rebound sleep.

Fig. 7 illustrates a simple model that assimilates our findings with those of Reichert et al. (2019): stimulant drugs drive increases in neuronal activity, as demonstrated by heightened *c-fos* expression, which drive a sleep pressure signal that is ultimately put into effect by release of galanin from the POA. This process can occur independently of noradrenalin-driven arousal. Given that noradrenergic signalling is a vital downstream effector for the arousing effects of hypocretin (Carter et al. 2012; Singh et al., 2015), the hypocretin system may also be dispensable for neuronal activity-induced rebound sleep, at least insofar as hypocretin-induced arousal relies on noradrenalin. This could be tested by performing stimulant-induced rebound sleep assays on hypocretin receptor knockout larvae.

**Fig. 7.**
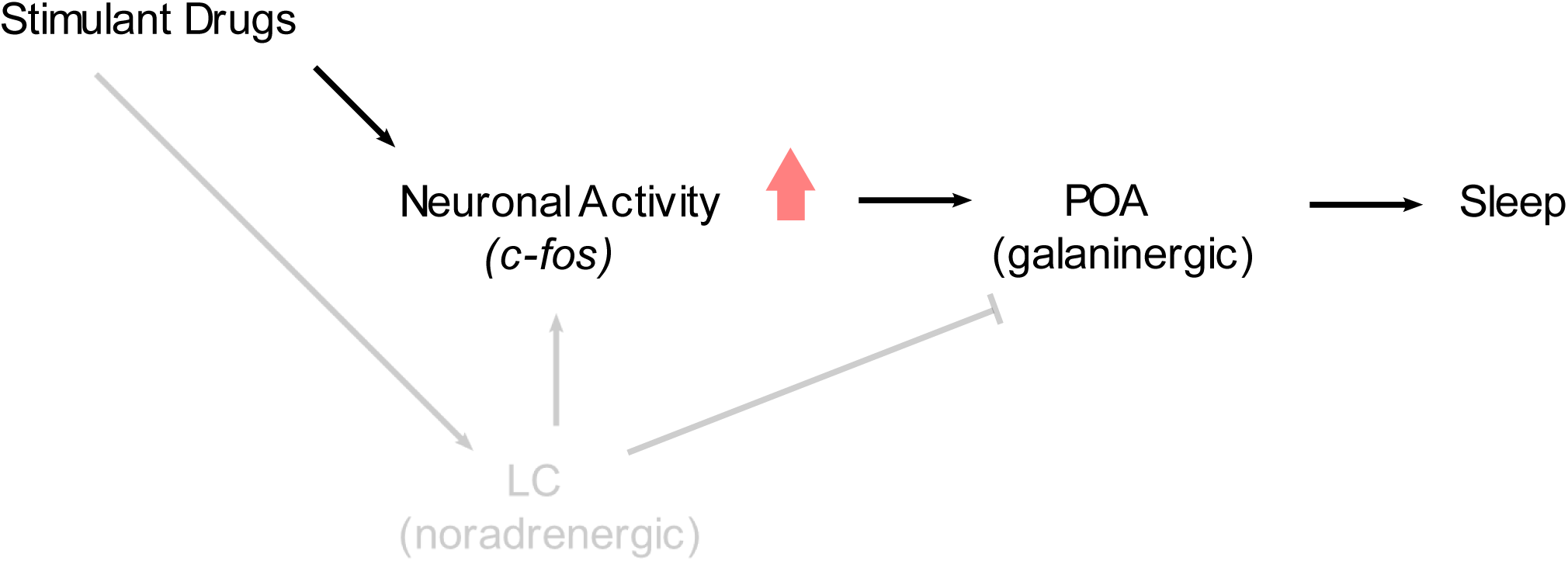
Noradrenergic activity is not required for stimulant-induced *c-fos* expression and rebound sleep. During waking, the LC releases noradrenalin to brain-wide targets, promoting arousal (Carter et al., 2010) and inhibiting sleep-promoting GABAergic/galaninergic neurons of the POA (Liang et al., 2021). Despite the role of the LC in maintaining arousal and heightened neuronal activity during waking, our results suggest that stimulant-induced neuronal activity and rebound sleep can occur in the absence of prior noradrenergic tone. Building on the work of Reichert et al. (2019), we propose a model in which stimulant-induced increases in neuronal activity subsequently promote activation of GABAergic/galaninergic sleep-promoting neurons of the POA, which drive sleep, independently of noradrenergic activity. Arrowheads denote activating projections; the bar head denotes an inhibitory projection

### A period of reduced noradrenergic activity could directly facilitate subsequent sleep

While an effect of magnified increases in neuronal activity on sleep pressure signalling is one plausible explanation of our results, another possibility is that the animal’s arousal state during waking directly affects subsequent sleep. When noradrenalin activates α_1_-adrenoceptors at excitatory glutamatergic synapses, this enhances synaptic transmission and can cause long-term potentiation (Perez 2020). Thus, reduced noradrenergic activity could relatively de-potentiate glutamatergic transmission in the wide-ranging brain regions to which the LC projects, limiting subsequent arousal. Cheng et al. (2020) suggest that in rats, sleep-promoting POA neurons receive excitatory glutamatergic afferents that promote sleep. Possible sources of these afferents include glutamatergic sleep-active neurons of the ventrolateral medulla, which reportedly directly excite POA GABAergic neurons in mice (Teng et al., 2022), and NREMS-promoting neurotensin-expressing glutamatergic neurons of the ventrolateral periaqueductal gray, which have been shown to project to the mouse POA (Zhong et al., 2019). Reduced noradrenergic activation of inhibitory α_2_-adrenoceptors on sleep-promoting POA neurons (Liang et al., 2021) might facilitate potentiation of these glutamatergic afferents (DeBock et al., 2003), thereby promoting sleep.

One seemingly paradoxical implication of direct inhibition of the sleep-promoting POA by noradrenalin is that α_2_-adrenoceptor agonists such as clonidine will also directly inhibit these sleep-promoting neurons. Indeed, McCarren et al. (2014) found that microinjection of the α_2_-adrenoceptor agonist dexmedetomidine into isoflurane-anaesthetised mouse ventrolateral POA increased behavioural arousal in vivo and reduced depolarisation in vitro. However, there is also evidence that noradrenergic inhibition of sleep-promoting neurons occurs indirectly, via activation of local GABAergic interneurons (Chamberlin et al., 2003; De Luca et al., 2022). α_2_-adrenoceptor agonists may therefore cause sedation when applied systemically because reduced noradrenergic activation of GABAergic interneurons that project to POA sleep-promoting neurons might outweigh the effects of the direct inhibition of the POA. This net disinhibition of sleep-promoting neurons would add to the general brain-wide sedating effects of α_2_-autoreceptor-mediated prevention of release of noradrenalin, along with the possible inhibitory heteroreceptor and postsynaptic effects of α_2_-adrenoceptor agonists.

The idea that noradrenergic activity during waking might affect subsequent sleep makes intuitive sense. To maximise survival, animals must optimally coordinate sleep and wake, balancing conflicting needs (Eban-Rothschild et al., 2018). A period of heightened noradrenergic tone might reflect an environmental change or threat, making sleep riskier than usual. A sleep-inhibiting after-effect of heightened noradrenergic arousal might therefore be adaptive. In *Drosophila*, Seidner et al. (2015) found that activating octopaminergic circuitry-the invertebrate counterpart of the noradrenergic system (Roeder 1999) - during sleep deprivation suppressed subsequent rebound sleep. One possible interpretation of this result is that sleep need continued to build during sleep deprivation, but counter-balancing after-effects of octopaminergic potentiation suppressed rebound sleep. Similarly, Suzuki et al. (2013) observed that mice kept awake by their spontaneous exploration of novel environments, which would engage the LC, showed greater sleep latencies afterwards than animals sleep deprived by gentle handling. Findings in other species are therefore at least consistent with the idea that changes in waking levels of noradrenergic/octopaminergic arousal can inversely impact subsequent sleep. To test the idea that noradrenergic after-effects on sleep occur due to plastic changes in synaptic transmission, experiments could be performed that measure electrophysiological changes in GABAergic/galaninergic POA neurons following opto-or chemo-genetic manipulation of the LC.

Nonetheless, our observation that clonidine boosts both baseline sleep and caffeine-induced rebound sleep in *dbh* F0 KOs is not consistent with the idea that clonidine enhances rebound sleep solely via the after-effects of its suppression of noradrenergic transmission. Rather, clonidine’s action on α_2_-adrenoceptors that sit on glutamatergic axon terminals, reducing the release of glutamate, and/or clonidine’s postsynaptic action as a neuronal inhibitor may also contribute to the rebound sleep enhancement that we observed. In any case, the interpretation that heightened immediate early gene expression explains the heightened rebound sleep in noradrenergic-compromised larvae does not preclude direct effects of prior noradrenergic tone on subsequent sleep; the two ideas are not mutually exclusive.

## Conclusion

Our results are consistent with previous findings in zebrafish that stimulant-induced rebound sleep increases as a function of preceding neuronal activity, as measured by *c-fos* expression. Additionally, we find that rebound sleep and *c-fos* expression are not dependent on heightened prior noradrenergic tone. In fact, reducing noradrenergic tone appears to *enhance* subsequent rebound sleep, perhaps by magnifying the increase in neuronal activity caused by the stimulant drug, as reflected by brain-wide levels of *c-fos* induction, and so augmenting a sleep pressure signal.

**Table 1.**
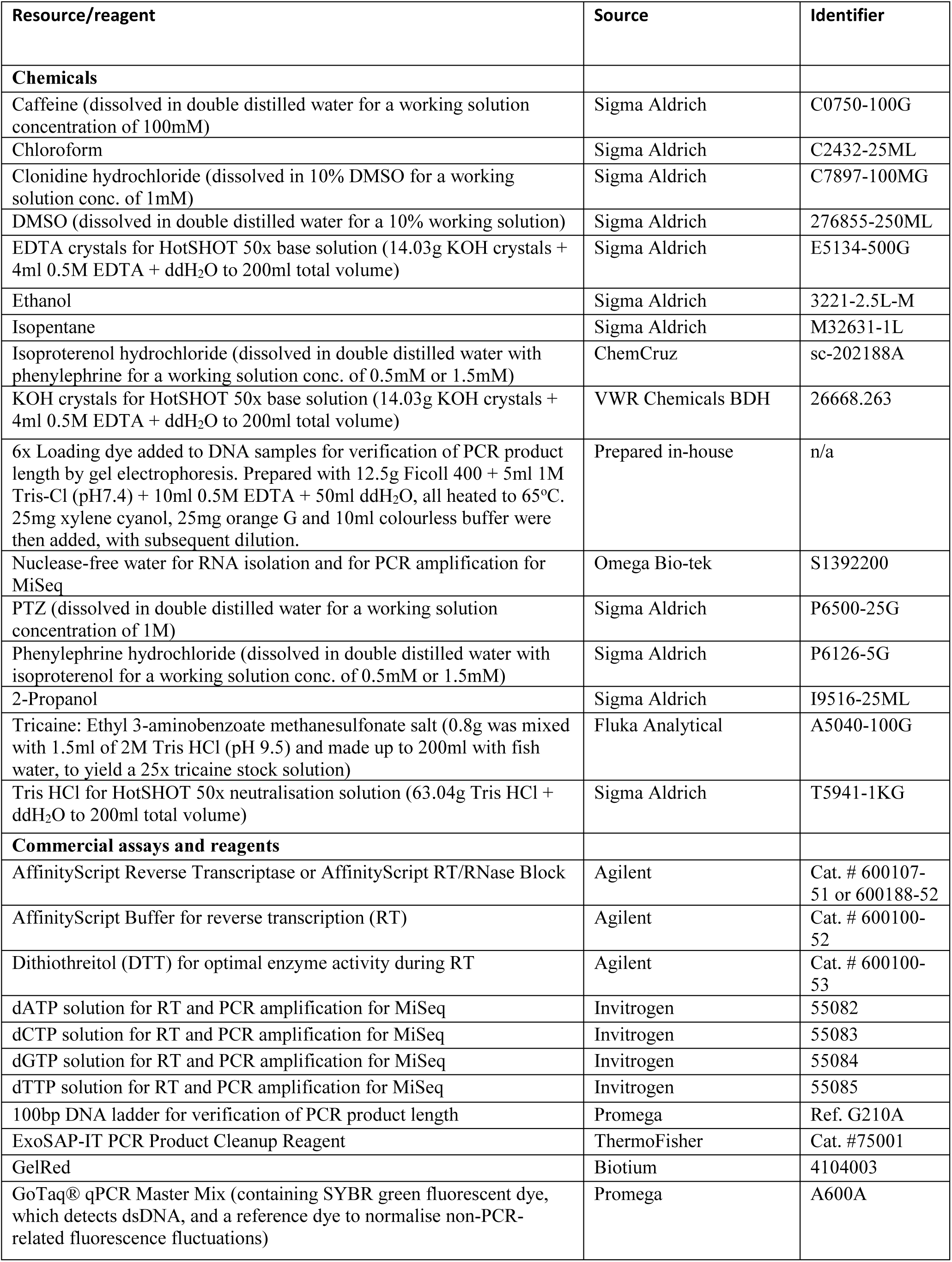
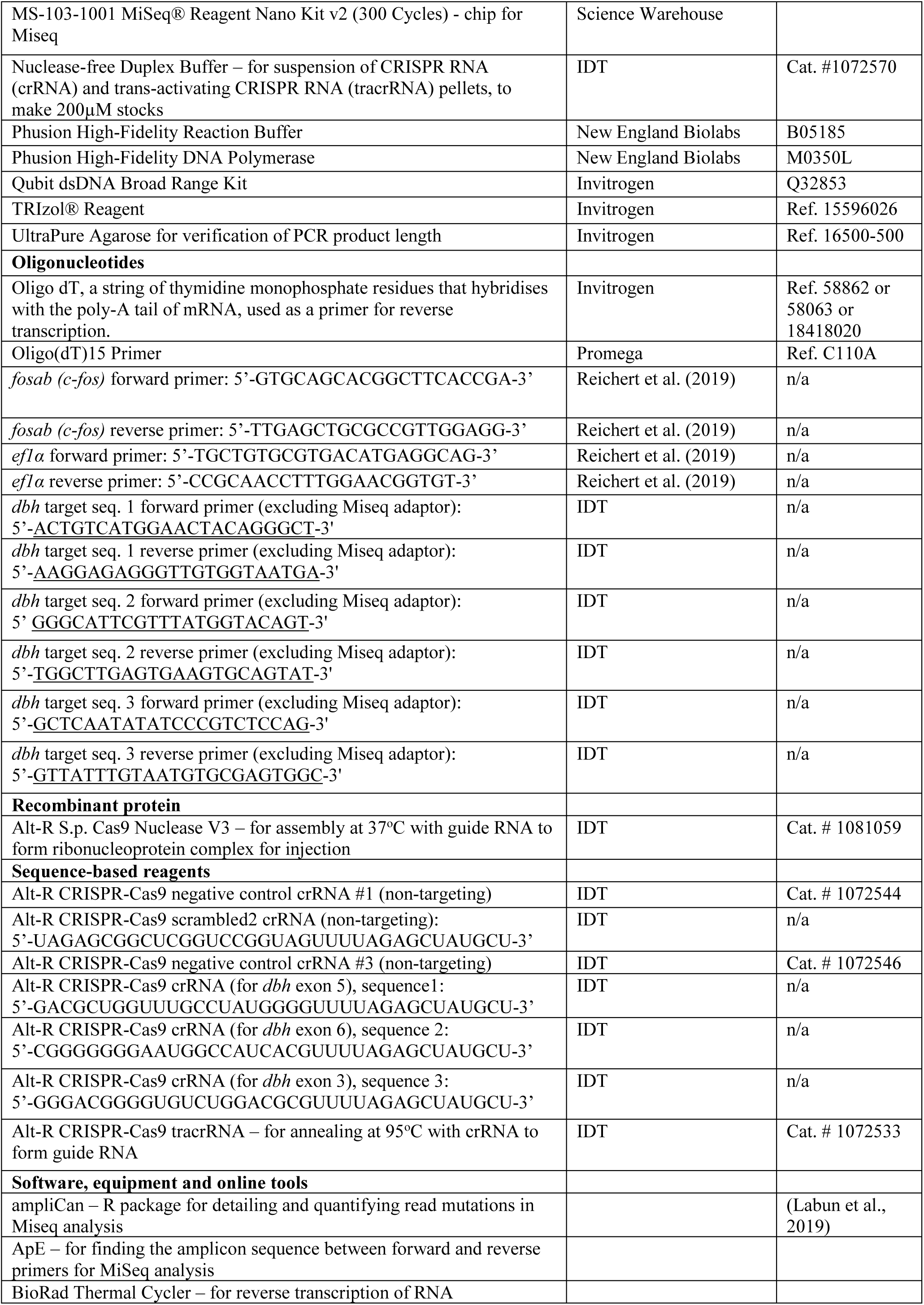
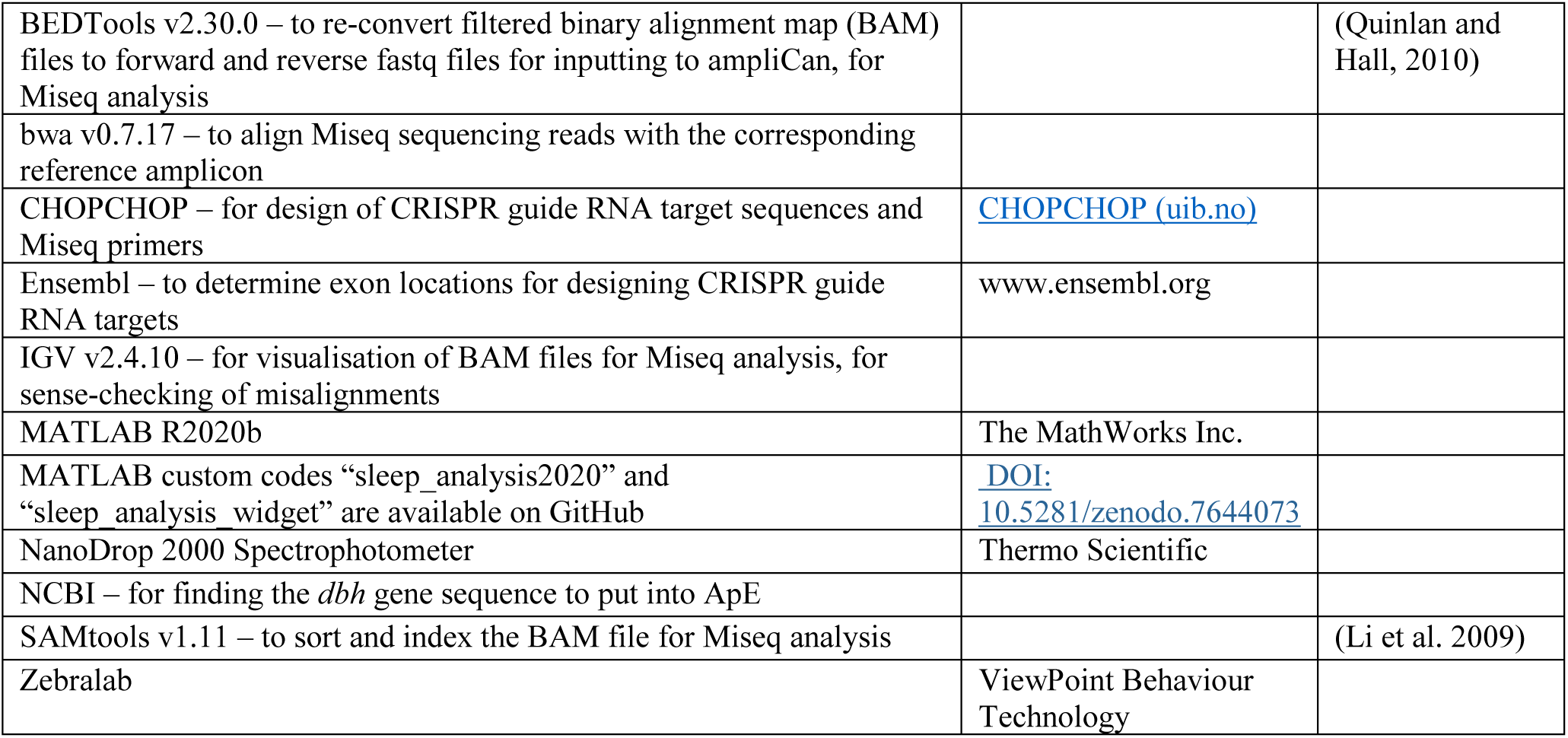
Key Resources

## Acknowledgements

David Prober kindly provided us with the underlying *dbh^-/-^* null mutant sleep data (Singh et al., 2015). We also thank the UCL Fish Facility for fish husbandry, Sumi Lim and Amanda Tan for qRT-PCR guidance, Francois Kroll for instruction in both designing targets for F0 knockout and in Miseq analysis, and all Rihel lab members for their ad hoc help and thoughts.

## Author contributions

All authors contributed to the conception and design of the study and interpretation of the results. EB performed the experiments, statistically analysed the data, prepared the figures and wrote the first draft of the manuscript. JR and DGL discussed and critically edited draft versions of the manuscript, and all authors read and approved the final version.

## Data availability

Data are available on request from the authors.

## Funding

This work was supported by an EMBO Fellowship awarded to DGL (ALTF 1097-2016), a European Research Council Starting Grant (282027) to JR, and a Wellcome Trust Investigator Award (217150/Z/19/Z) to JR.

## Declarations

### Conflicts of interest

The authors declare that they have no conflicts of interest.

### Ethical approval

Experiments and zebrafish husbandry followed UCL fish facility protocols per project licence PA8D4D0E5, awarded to Jason Rihel by the UK Home Office under the UK Animals (Scientific Procedures) Act 1986.

---

**Fig. S1.**
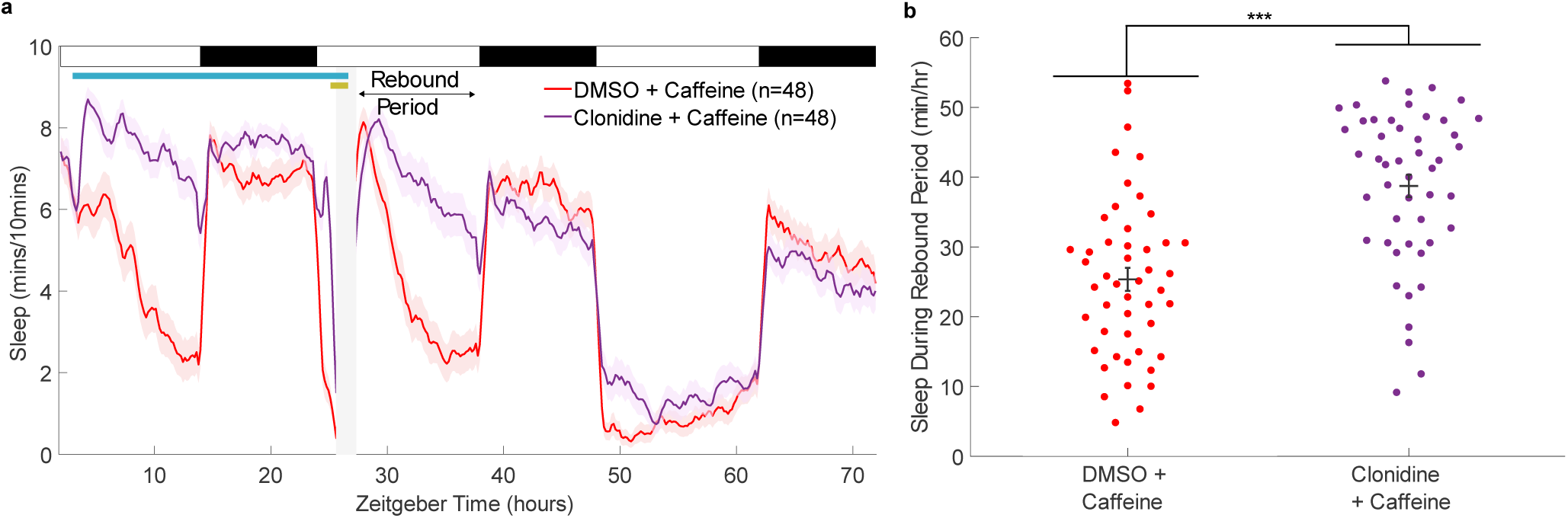
Stimulation of α2-adrenoceptors during caffeine-induced arousal increases rebound sleep. **a** Sleep traces (± SEM) beginning at 5 dpf and continuing over three days and nights (ZT0 = first lights on) for larvae exposed to either DMSO+caffeine or clonidine+caffeine. Following drug wash-off, rebound sleep is enhanced by clonidine. At top left, the pale blue horizontal bar shows the clonidine exposure window while the gold bar indicates the presence of stimulant. **b** shows the average total sleep/hr during the rebound sleep period for each larva (black cross: mean ± SEM). Rebound sleep was significantly higher following treatment with clonidine than treatment with vehicle (p=8.8 x 10^-8^, F(1,94) = 33.67), one-way ANOVA. ***p<0.001

**Fig. S2.**
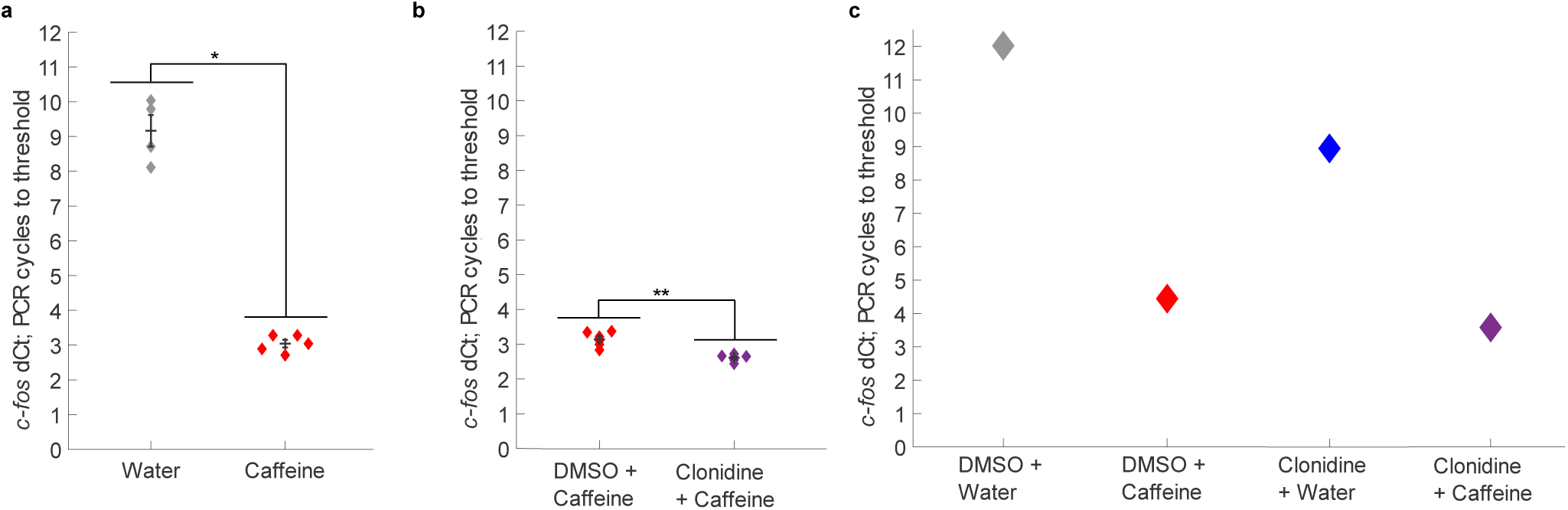
*c-fos* expression is greater in larvae exposed to both clonidine and caffeine than in larvae exposed to only caffeine. **a** Larvae treated with water (n = 4 groups of ∼20 larvae) required significantly more PCR cycles for *c-fos* cDNA amplification to achieve threshold fluorescence, normalised to *ef1α* cycles-to-threshold, than larvae treated with caffeine (n = 5 groups of ∼20 larvae); W’ = 10, p = 0.0159, two-tailed Wilcoxon rank sum test (black cross: mean ± SEM). **b** Larvae treated with vehicle and caffeine (n = 6 groups of ∼20 larvae) required significantly more PCR cycles for *c-fos* cDNA amplification to achieve threshold fluorescence, normalised to *ef1α* cycles-to-threshold, than larvae treated with clonidine and caffeine (n = 6 groups of ∼20 larvae); W = 21, p = 0.0022, two-tailed Wilcoxon rank sum test. **c** Each datapoint represents 1 group of 37 larvae. The number of normalised *c-fos* PCR cycles to threshold was highest in the vehicle-only condition and lowest in the clonidine + caffeine condition. *p<0.05, **p<0.01. Each datapoint is the mean of three technical replicates

**Fig. S3.**
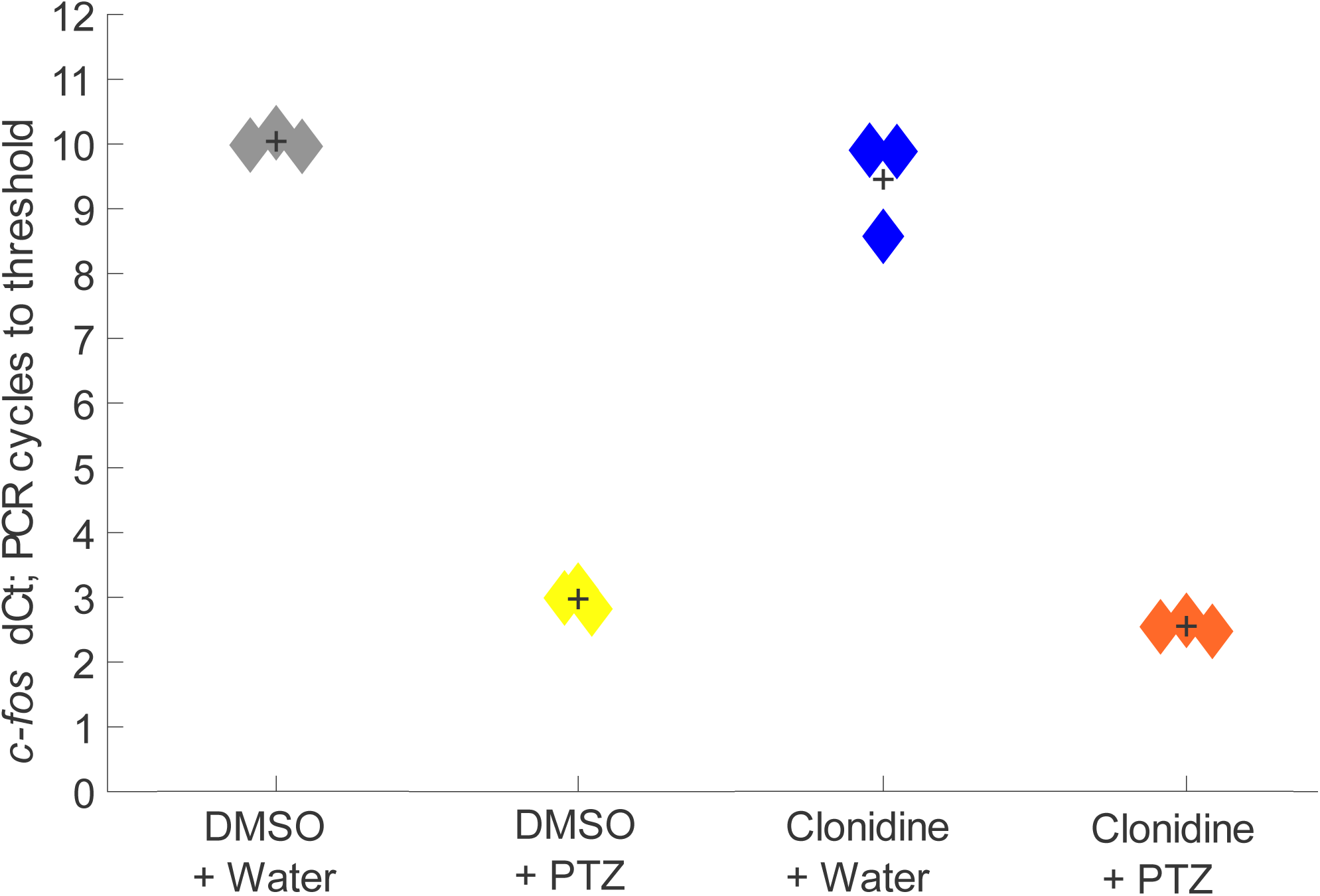
*c-fos* PCR cycles to threshold across different clonidine/PTZ treatment combinations. qRT-PCR on groups of ∼17 larvae (n=3 biological replicates per condition) reveals that the average number of normalised *c-fos* PCR cycles to threshold is highest in the vehicle-only condition and lowest in the clonidine + PTZ condition. Black cross: mean. Each datapoint is the mean of three technical replicates

**Fig. S4.**
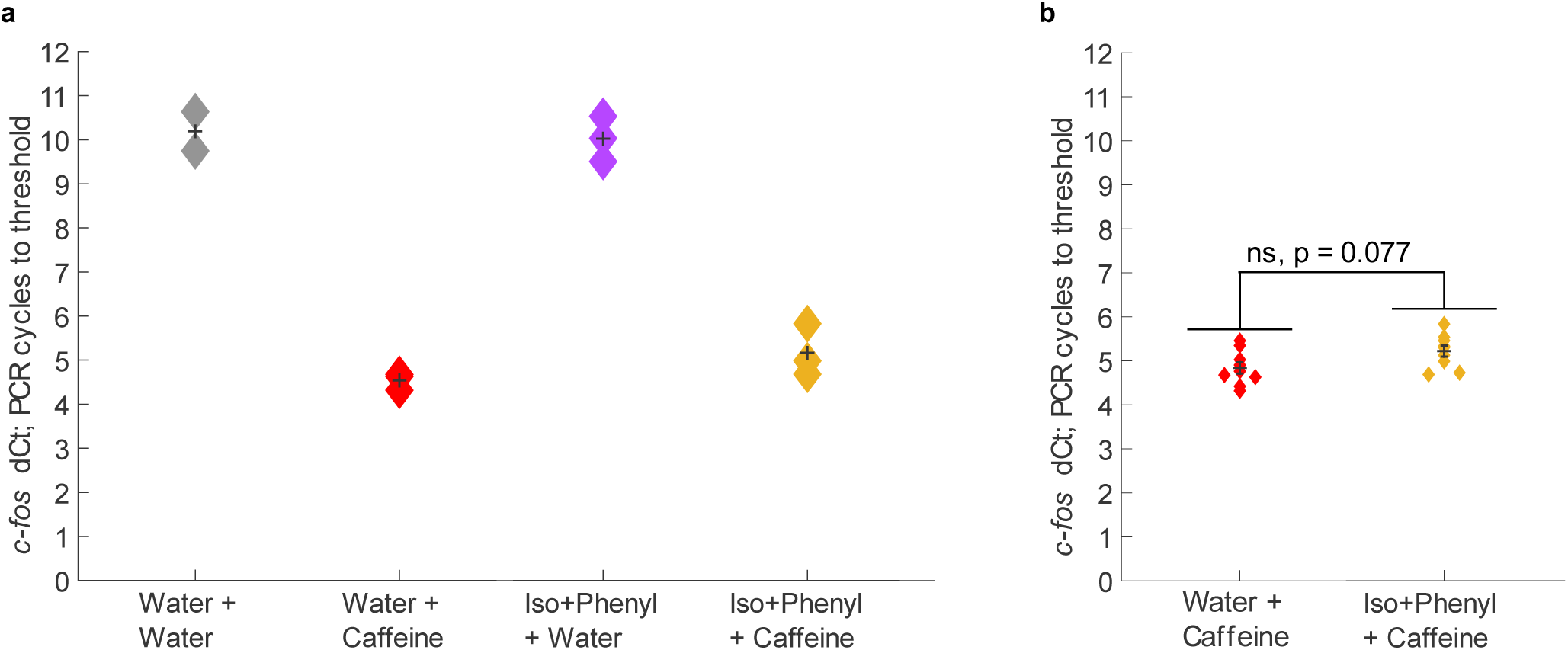
*c-fos* PCR cycles to threshold across different isoproterenol+phenylephrine/caffeine treatment combinations. **a** qRT-PCR on groups of ∼18 larvae (n=3 or n=2 biological replicates per condition) reveals that the average number of normalised *c-fos* PCR cycles to threshold is lower in the caffeine condition and higher in the isoproterenol+phenylephrine and caffeine condition. **b** includes the three water and caffeine datapoints and the three isoproterenol+phenylephrine and caffeine datapoints from **a**. An additional 6 groups of ∼20 larvae were treated with water and caffeine while 6 groups were treated with isoproterenol+phenylephrine and caffeine, and qRT-PCR analysis was conducted. Larvae treated with water and caffeine (n = 9 biological replicates) did not require significantly fewer PCR cycles for *c-fos* cDNA amplification to achieve threshold fluorescence, normalised to *ef1α* cycles-to-threshold, than larvae treated with isoproterenol+phenylephrine and caffeine (n = 9 biological replicates); W = 65, p = 0.077, two-tailed Wilcoxon rank sum test. Each datapoint is the mean of three technical replicates

**Fig. S5.**
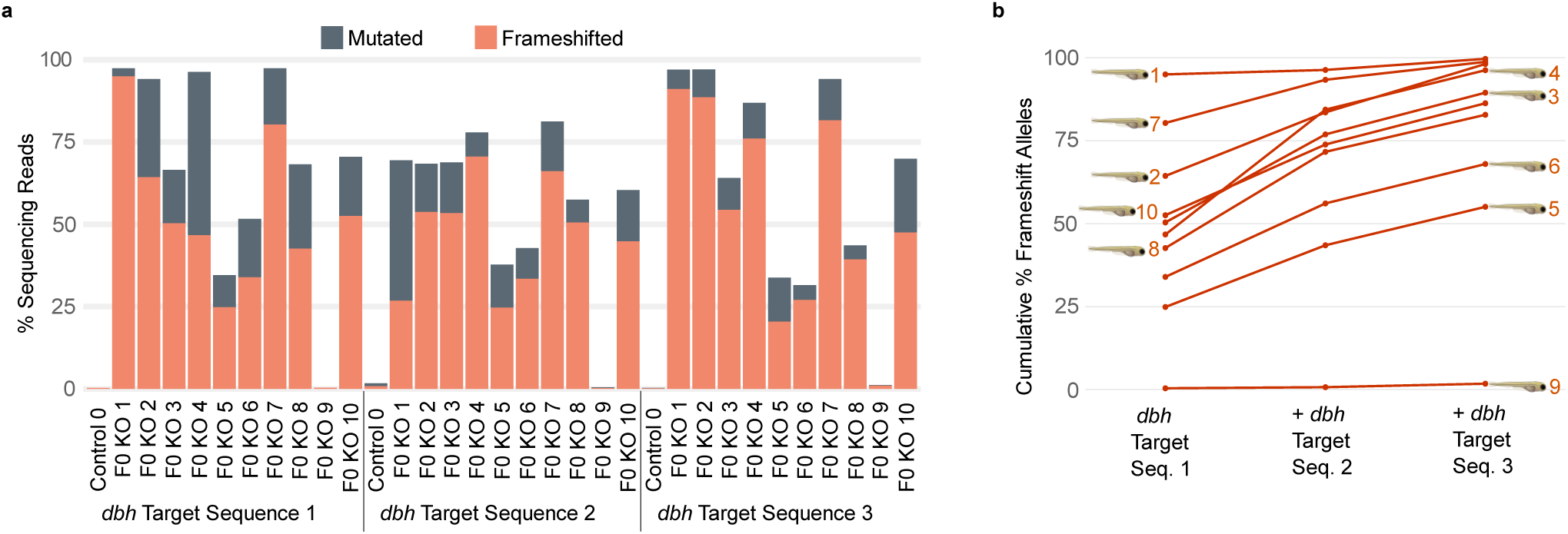
Deep sequencing reveals a high proportion of frameshifted *dbh* copies among most *dbh* F0 KO larvae, indicative of highly penetrant, largely null mutations. **a** shows the percentage of reads with a mutation of any kind (full heights of the bars) and the percentage of reads with a frameshift mutation (orange portion) for DNA samples from 10 blindly chosen *dbh* F0 KO larvae, and one control-injected larva, that were included in the experiment shown in Fig. S6. For each larva, mutation counts are shown for each of the 3 sequences within the *dbh* gene that were targeted by the CRISPR/Cas9 injections. **b** illustrates the cumulative proportion of *dbh* copies in each F0 KO larva that are estimated to have a frameshift mutation, considering all target sequences together. Each of the 10 orange lines represents one *dbh* F0 KO larva, numbered as per **a**

**Fig. S6.**
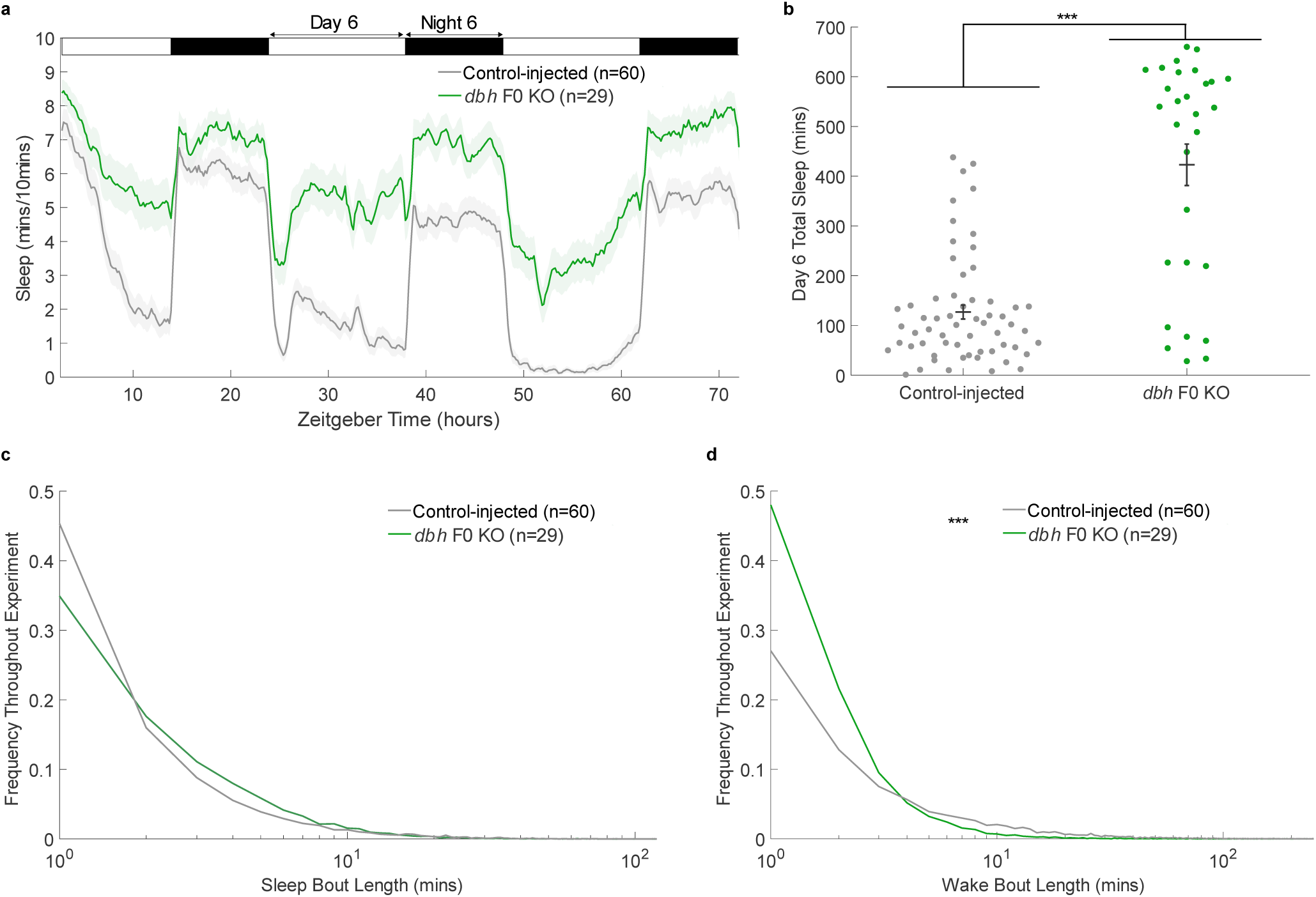
d*b*h F0 KO larvae show significantly higher daytime sleep levels than controls. **a** Sleep traces for two groups of zebrafish larvae that had been injected at the 1-cell stage with CRISPR/Cas9 guide RNAs. “*dbh* F0 KO” larvae were injected with guide RNAs that targeted the *dbh* gene. Control-injected larvae were injected with guide RNAs whose sequences are not predicted to match any genomic region. Day 6 total sleep is summarised for each larva in **b**. *dbh* F0 KO larvae showed significantly higher day 6 total sleep than control-injected larvae; p=7.8 x 10^-13^, F(1,87) = 70.47, one-way ANOVA. **c** shows the distribution of sleep bout lengths of *dbh* F0 KO and control-injected larvae over the course of the tracking experiment. *dbh* F0 KO larvae had fewer short sleep bouts and more long sleep bouts than controls, though the effect was not statistically significant (p>0.05, Kolmogorov-Smirnov test). **d** illustrates that *dbh* F0 KO larvae had significantly more short wake bouts and fewer long wake bouts than controls (p=1.3 x 10^-7^, Kolmogorov-Smirnov test). ***p<0.001

**Fig. S7.**
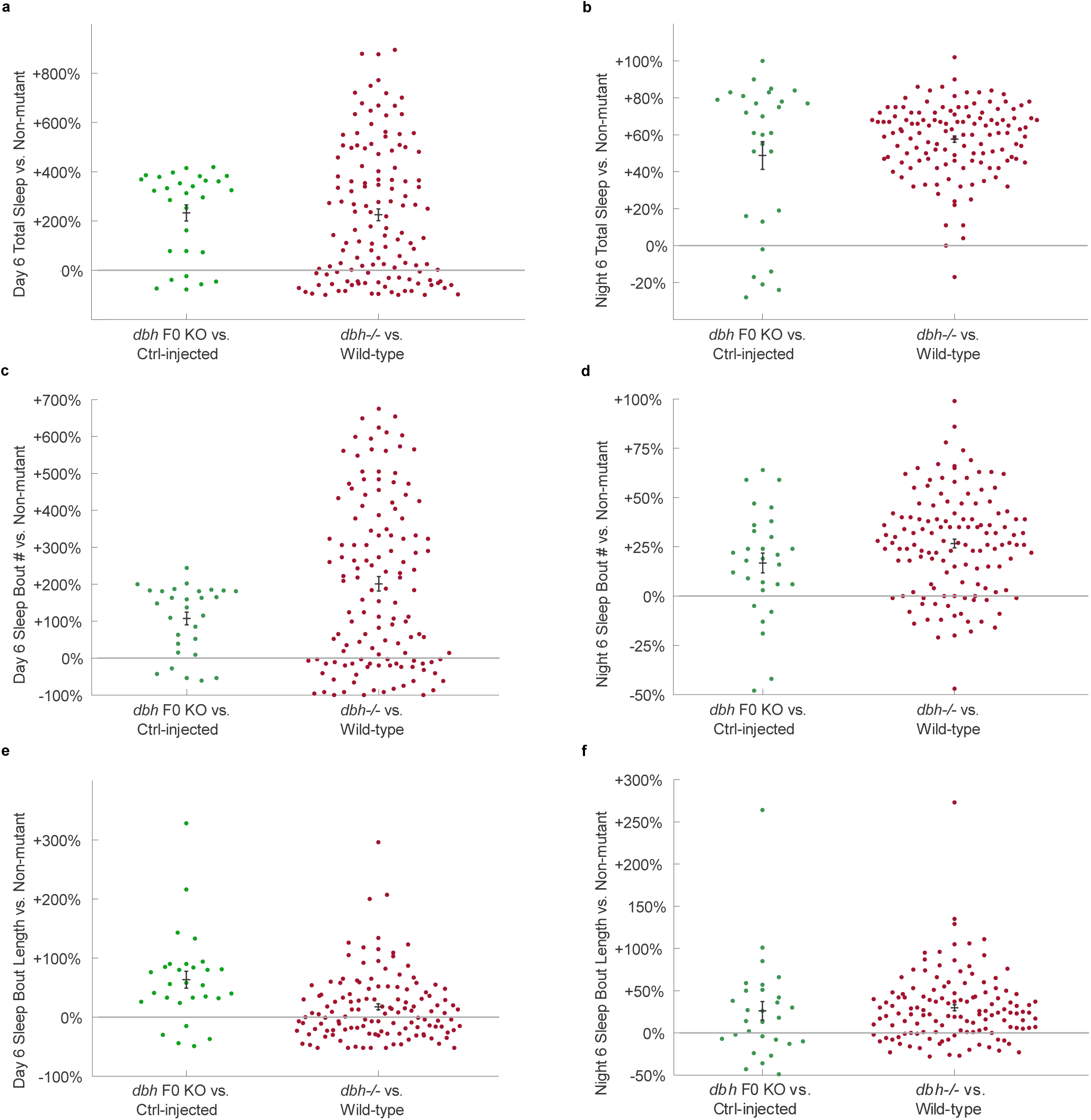
d*b*h F0 KOs show a similar sleep phenotype to previously described *dbh*^-/-^ null mutants. **a** shows the day 6 total sleep levels of *dbh* F0 KO larvae compared with control-injected larvae alongside the day 6 total sleep levels of *dbh^-/-^* null mutants compared with wild-type larvae; **b** compares night 6 total sleep levels. **c** and **d** compare sleep bout numbers during day 6 and night 6, respectively and **e** and **f** compare sleep bout lengths during day 6 and night 6. The underlying data for *dbh* F0 KO and control-injected larvae are from the experiment in Fig. S6. The data for *dbh^-/-^* null mutants and wild-type larvae were provided by David Prober, as reported in Singh et al. (2015). Each dot represents the data for one F0 KO or mutant larva, normalised to the mean value for all control-injected or all wild-type larvae, respectively

**Fig. S8.**
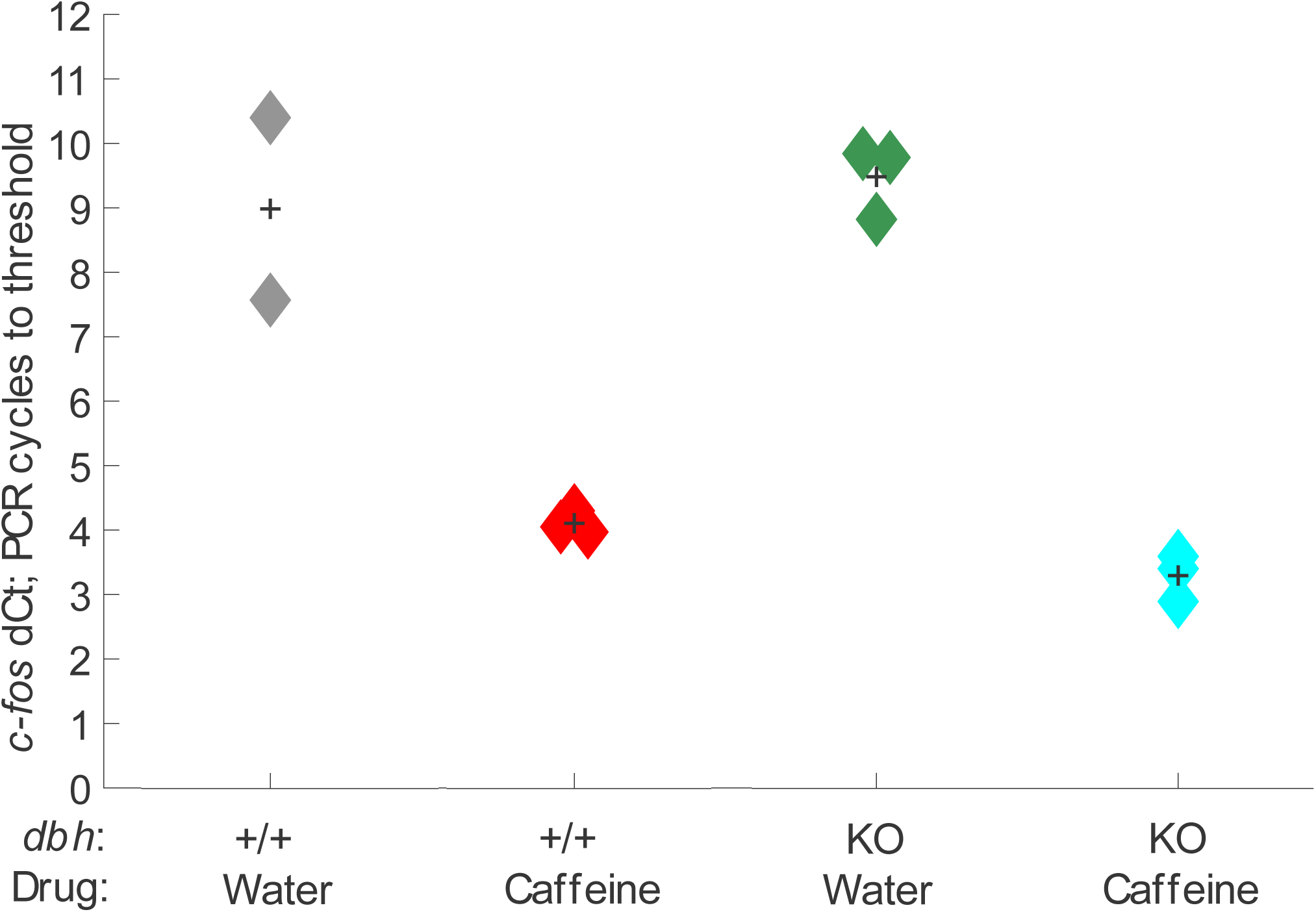
*c-fos* PCR cycles to threshold across groups of *dbh* F0 KOs and control-injected larvae treated with caffeine or water. qRT-PCR on groups of ∼15 larvae reveals that the number of normalised *c-fos* PCR cycles to threshold for each group of *dbh* F0 KO larvae treated with caffeine (n=3 biological replicates) was lower than for the groups of control-injected larvae treated with caffeine (n=3 biological replicates). Each datapoint is the mean of three technical replicates

**Fig. S9.**
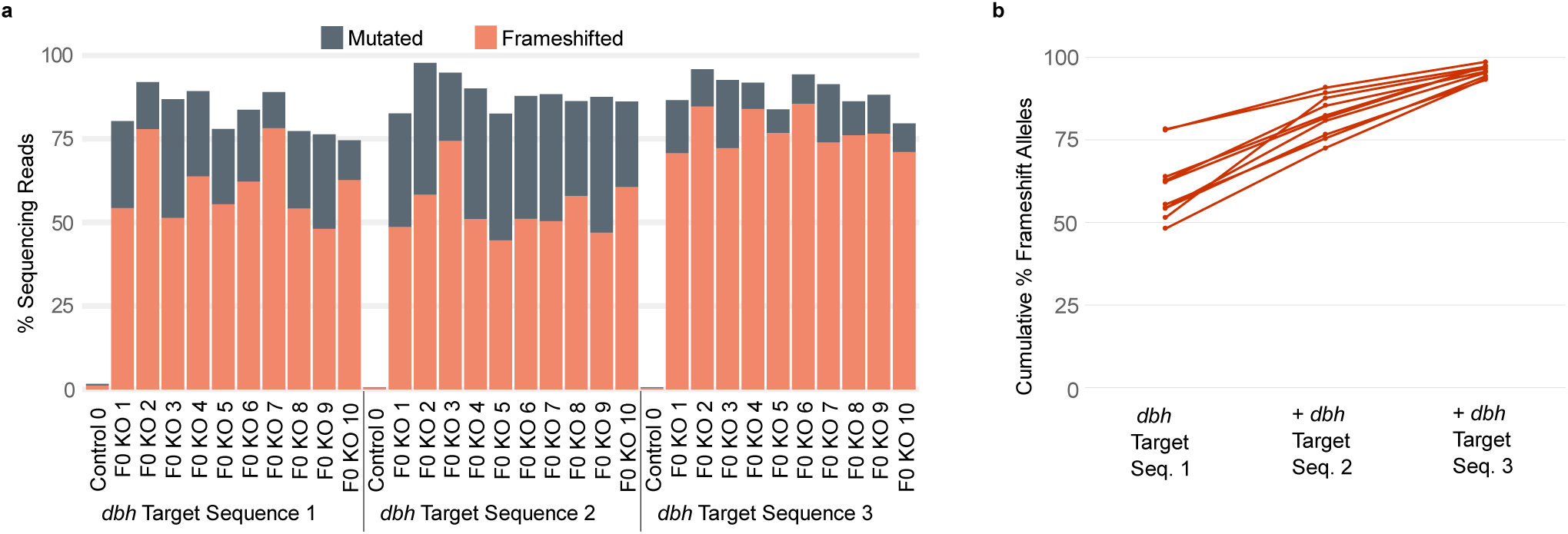
Deep sequencing reveals a high proportion of frameshifted *dbh* copies among *dbh* F0 KO larvae. **a** reveals that the majority of reads from DNA samples of 10 *dbh* F0 KO larvae (Fig. 7) have frameshift mutations. **b** shows the cumulative proportion of *dbh* copies in each F0 KO larva that are estimated to have a frameshift mutation, considering all target sequences together. All larvae had a cumulative frameshift rate of >93% (mean 96%), indicating a high penetrance of loss-of-function mutations in the *dbh* F0 KOs of the experiment in Fig. 7

## Notes

### Competing Interest Statement

The authors have declared no competing interest.

